# Centromere landscapes resolved from hundreds of human genomes

**DOI:** 10.1101/2024.01.26.577337

**Authors:** Shenghan Gao, Yimeng Zhang, Stephen J. Bush, Bo Wang, Xiaofei Yang, Kai Ye

## Abstract

High-fidelity (HiFi) sequencing has facilitated the assembly and analysis of the most repetitive region of the genome, the centromere. Nevertheless, our current understanding of human centromeres draws from a relatively small number of telomere-to-telomere assemblies, and so has not yet captured its full diversity. In this study, we investigated the genomic diversity of human centromere higher order repeats (HORs) using both HiFi reads and haplotype-resolved assemblies from hundreds of samples drawn from ongoing pangenome-sequencing projects and reprocessed using a novel HOR annotation pipeline, HiCAT-human. We use this wealth of data to provide a global survey of the centromeric HOR landscape, in particular finding that 23 HORs exhibited significant copy number variability between populations. We detected three centromere genotypes with imbalance population frequencies on each of chromosome 5, 8 and 17. An inter-assembly comparison of HOR loci further revealed that while HOR array structures are diverse, they nevertheless tend to form a number of specific landscapes, each exhibiting different levels of HOR subunit expansion and possibly reflecting a cyclical evolutionary transition from homogeneous to nested structures and back.

## Introduction

Centromeres are essential, yet rapidly-evolving, chromosomal domains with functional roles in cell division ^1^, although are characteristically challenging to assemble ^2^. Human centromere sequences typically comprise multiple alpha satellite monomers (of length ∼171bp, and generally sharing 50%-90% identity) organized into higher order repeat (HOR) units (which share approx. 95-100% identity) ^3,4^.

Such a high level of repetition ensures that centromeres are difficult to assemble and that reads cannot easily be mapped to them with high accuracy, collectively hindering investigations of centromere architecture and evolution ^4,5^. However, advanced long read sequencing technologies, in particular PacBio high-fidelity (HiFi) reads^6^, have recently achieved complete centromere assembly, with the Telomere-to-Telomere (T2T) consortium publishing the first complete human genome (CHM13) in 2022 alongside an analysis of its centromeres^2,7^. This provided a detailed chromosome-specific HOR atlas and demonstrated that human centromeres evolve by a process of “layered expansion” in which younger sequences expand from the middle, in a manner resembling successive tandem duplications, with older flanking sequences shrinking and diverging over time^7^. A second T2T human genome has since been completed (CHM1) alongside analyses of both the genetic and epigenetic variation within its centromeres ^8^.

Despite the substantial insight afforded by assembling genomes to T2T level, in absolute terms a small number of complete assemblies remains insufficient for characterizing the rapid evolution and diversity of centromere sequence. To that end, Suzuki *et al.* used long-read sequencing of 36 individuals to reveal the structural diversity of human centromere HORs ^9^. However, their samples primarily comprised individuals from one population (Japanese) and their strategy was limited to characterising the HOR patterns of chromosomes 11 (chr11), 17 and X (collectively ‘suprachromosomal family 3’) as these were considered more divergent other chromosomes ^9,10^. Recently, both the Human Pangenome Reference Consortium (HPRC) and the Chinese Pangenome Consortium (CPC) have released HiFi reads and haplotype-resolved assemblies of over a hundred individuals from a diverse range of ancestries, which to the best of our knowledge have not yet been used in a dedicated population-wide analysis of centromere sequence variation ^11,12^. Combining data from these projects provides an opportunity to investigate the diversity and evolution of centromere sequences among the broader human population. Here, we refine and update our previous HOR-annotation tool, HiCAT^13^, initially designed only for use with individual assembly, to annotate hundreds of centromeres. Using this improved version, HiCAT-human, we analysed HiFi reads from 102 individuals and the assemblies of 109 haplotypes – collectively, every sample released by both the HPRC and CPC, plus CHM13 – leveraging this wealth of data to provide a comprehensive global survey of the human centromeric landscape.

## Results

### HOR annotation in human population

To investigate the diversity of human centromeres, we analysed HPRC and CPC HiFi reads from 102 individuals including 22 from Africa (AFR; all from HPRC), 16 from Latin America (AMR, all from HPRC), 62 from East Asia (EAS, 4 from HPRC and 58 from CPC), and 1 from South Asia (SAS, from HPRC), plus CHM13. In addition, we analysed 108 haplotype-resolved assemblies (43×2 from HPRC and 11×2 from CPC) alongside the CHM13 genome (Supplementary table S1) ^2,11,12^.

We modified our HOR annotation tool HiCAT, which was originally designed to work with individual assemblies ^13^, to create an updated version, HiCAT-human, which can automatically annotate centromere HOR patterns from both reads and assemblies of multiple human samples (Fig. 1, Supplementary figure S1, Supplementary table S2, S3 and Methods). HiCAT-human comprises two workflows for this purpose, HiCAT-human-reads and HiCAT-human-assembly. In HiCAT-human-reads, alpha satellite reads (ASR) are extracted and classified into each chromosome with a pre-trained classifier (Fig. 1a), and in HiCAT-human-assembly, alpha satellite regions in each chromosome are detected using Lastz ^14^ and then merged to obtain a chromosome-specific alpha satellite array (CASA) (Fig. 1b). In the subsequent annotation step, both ASRs and CASAs were transformed into block sequences using StringDecomposer ^15^ and then into monomer sequences using monomer templates derived by HiCAT ^13^. Monomer sequences were used to derive HORs following a hierarchical tandem repeat mining (HTRM) approach ^13^ (Fig. 1c).

**Fig. 1.**
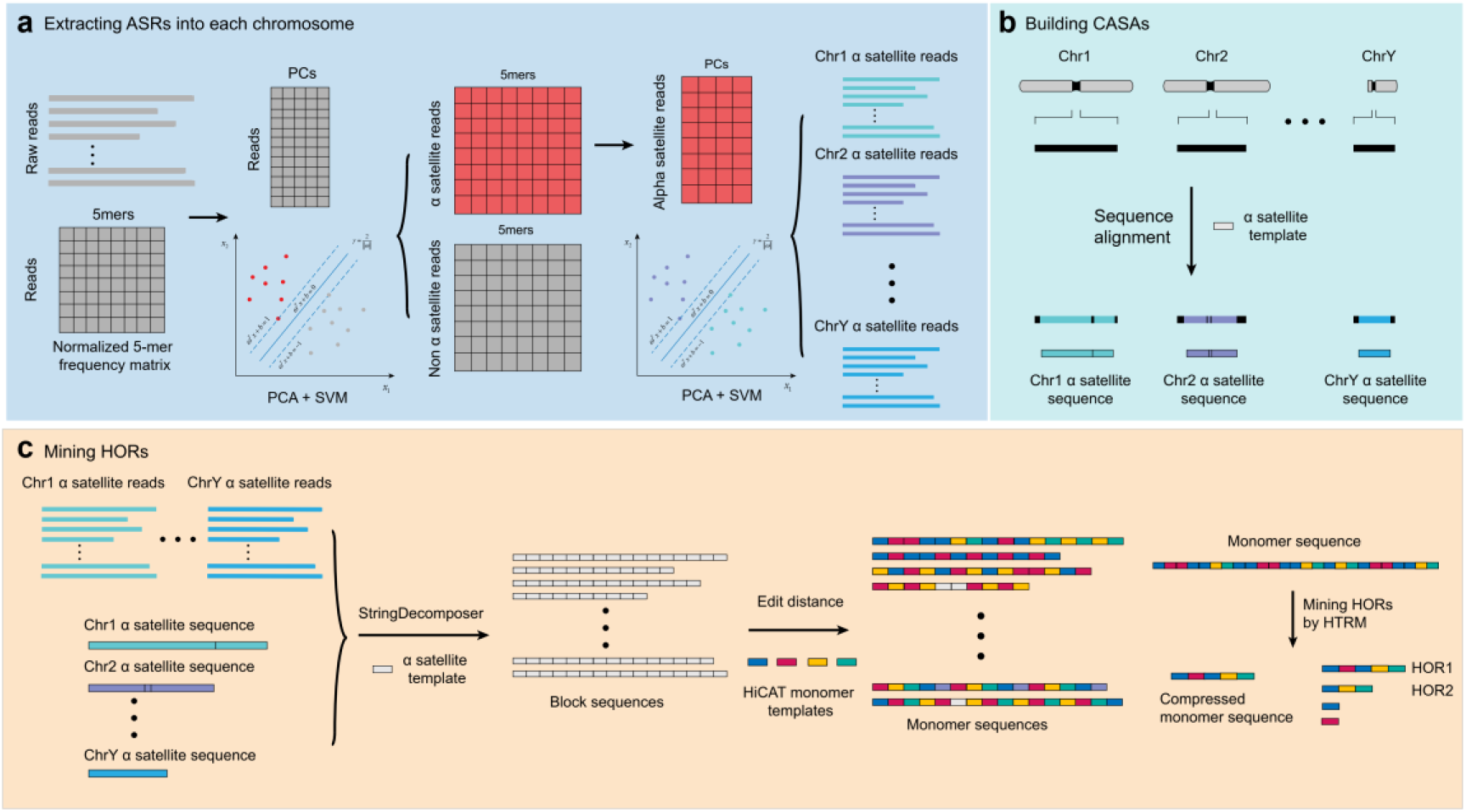
Overview of HiCAT-human. **a.** Extracting reads containing alpha satellite and classifying them into each chromosome. 5-mer frequency matrix constructed from raw reads (grey) are classified into alpha satellite reads and non-alpha satellite reads using principal component analysis (PCA) and support vector machine (SVM). Then, 5-mer frequency matrix constructed from alpha satellite reads (red) are classified into each chromosome using PCA and SVM. **b.** Building chromosome specific alpha satellite array (CASA) from assembled genome. Black bars represent pericentromeric and centromeric regions. Regions inside bars with different colors represent alpha satellite regions in each chromosome. Sequence alignment is performed by Lastz ^14^ using alpha satellite as template. **c.** Mining higher order repeats (HORs). All chromosomes’ ASRs and CASAs are first transformed into block sequences based on StringDecomposer ^15^ with the alpha satellite sequence as a template, with the edit distance between the HiCAT monomer templates and blocks used to transform block sequences into monomers. Finally, HORs were annotated using the monomer sequences and the hierarchical tandem repeat mining (HTRM) method ^13^. Different coloured rectangles in the monomer sequence represent different monomers.

### Human HOR quantification based on HiFi reads data

To identify human centromere HORs, we applied the HiCAT-human-reads workflow to the HiFi reads of 102 samples. We estimated the size of the HOR arrays for each chromosome in each sample as the total number of bases in the HOR reads divided by the sequencing coverage. The median HOR array size varied from 0.8 to 4.5 Mb across all chromosomes (Fig. 2a and Supplementary figure S2a) and showed marked variability between populations. We found that for eight chromosomes, HOR arrays in EAS populations were significantly larger than those of AFR and AMR and that conversely, in chr16, 21 and Y, the arrays were significantly larger in AFR populations than others (Supplementary figure S3-5, Supplementary table S4). To compare how the number of HORs varied among samples, we calculated the ‘n-number’ as the total count of HORs in each sample normalized by depth of sequencing coverage (Methods). For subsequent analysis, we excluded rare HORs to capture the main characteristics and avoid artefacts (Methods). In total, we obtained 79 HORs, 33 of which exhibited pronounced variance between populations (on the basis of a mean fold-change in their n-number among samples; Methods), and which we refer to as ‘variable HORs’ (v-HORs) (Fig. 2b and Supplementary table S5-S11). This variation in normalised HOR counts demonstrates that the CHM13 genome represents only one distribution of the human HOR landscape and that by extension it is unable to capture the broader range of human HOR diversity (Fig. 2b and Supplementary figure S2b, c). We found that 23 of the 33 v-HORs were significantly variable among populations (Fig. 2c, Supplementary figure S6, S7 and Supplementary table S12), including 5_M2L8 (significantly higher in AMR than in EAS) and 8_M4L7 (significantly higher in EAS than that all other populations analysed).

**Fig. 2.**
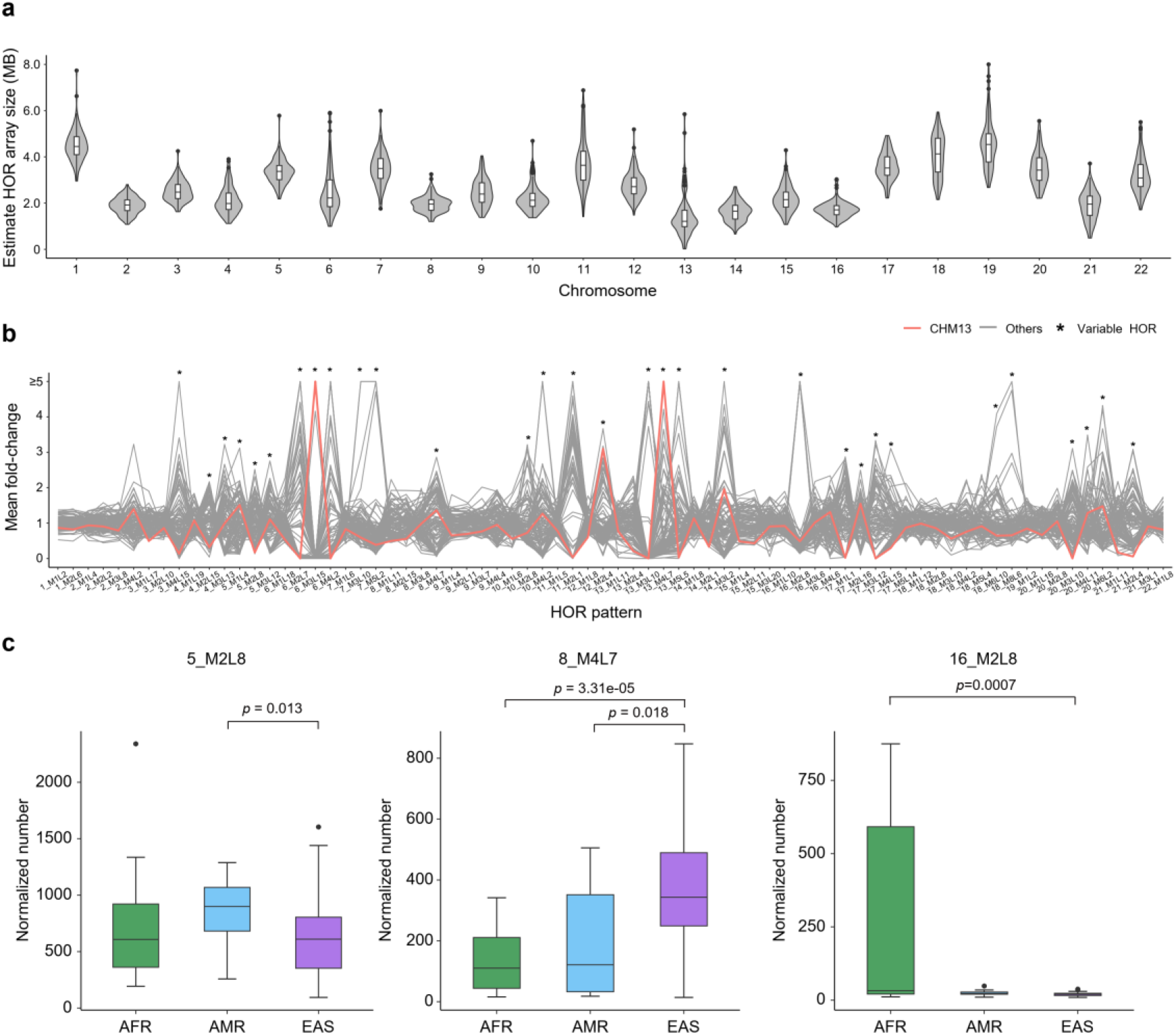
HOR annotation for HiFi reads of 102 individuals. **a.** The estimated HOR array size based on total length of HOR reads and sequencing coverage in each autosome. **b.** The mean fold-change of autosomal HORs among all samples. Mean fold-change is calculated by dividing the normalized numbers (n-number) of each HOR in each sample by the mean n-number of that HOR across all samples. CHM13 is represented by a red line and all other samples by grey lines. v-HORs are marked by asterisks. The results for sex chromosomes are shown in Supplementary figure S2**. c.** The n-number of three representative HOR arrays (5_M2L8, 8_M4L7 and 16_M2L8) in AFR (sample number, n = 22), AMR (n = 16) and EAS (n = 62) populations. Other v-HORs are shown in Supplementary figure S6 and S7. *p*-value is calculated by two-sided Wilcoxon rank sum test.

### Chromosomal variability in centromere genotypes

To explore the distribution of HORs between chromosomes, we calculated the correlation between the n-number of different HORs in all samples and consistently found no significant correlation between inter-chromosomal HORs and a high correlation between intra-chromosomal HORs (Supplementary figure S8 and Supplementary table S13), which suggests very little commonality between chromosomes in terms of their centromere composition. For each of the 17 chromosomes with v-HORs (i.e., those chromosomes whose HORs show pronounced variation between samples), samples could be grouped generally into two or three clusters (Fig. 3, Supplementary figure S9-17 and Supplementary table S14). Taking chr5 as an example, we detected three clusters (termed 5_C0, 5_C1 and 5_C2), each with a variable composition of HORs (Fig. 3a, b and Supplementary figure S9a). We found that the n-numbers of all three HORs in cluster 5_C2 approximated the averaged n-number of clusters 5_C0 and 5_C1 (Fig. 3a, c and Supplementary table S15), which suggests that the latter are homozygous (AA and BB, respectively) and that by extension cluster 5_C2 is heterozygous (AB). Moreover, we found that 5_C2 was more frequently detected in each of the three main population groups (being present in 36-56% of samples) whereas 5_C0 (AA) and 5_C1 (BB) showed greater population bias (Fig. 3d). Specifically, the EAS population had the significantly lowest proportion of samples with the 5_C0 (AA) genotype (12.9%) (*p*-value is 0.039 compared to AFR and 1.636e-05 compared to AMR, one-sided binomial test) while the AMR population had the significantly lowest proportion of 5_C1 (BB) (6.3%, *p*-value is 0.003 compared to AFR and 0.017 compared to EAS; Fig. 3d). We found similar results for three clusters of HORs on chr8 (Fig. 3e-g and Supplementary table S15) and chr17 (Supplementary figure S17 and Supplementary table S15). In chr8, we found that no 8_C1 (AA) is in AFR samples and the sample ratios of 8_C0 in AMR (50.0%, *p*-value is 0.002) and AFR (36.4%, *p*-value is 0.017) are significantly higher than that of EAS (16.1%) (Fig. 3h). In chr17, the three genotypes were reported in previous studies with allele frequency of B is 61.9% in Japanese population ^9^ and 35% in European population ^16^. We found that allele frequency of B in EAS is 0.476 (59 of 124) which is significantly higher than European population (*p*-value is 0.0026) but lower than Japanese population (*p*-value is 0.0008). Taken together, these results suggest that centromeric genotypes are highly variable among and between populations.

**Fig. 3.**
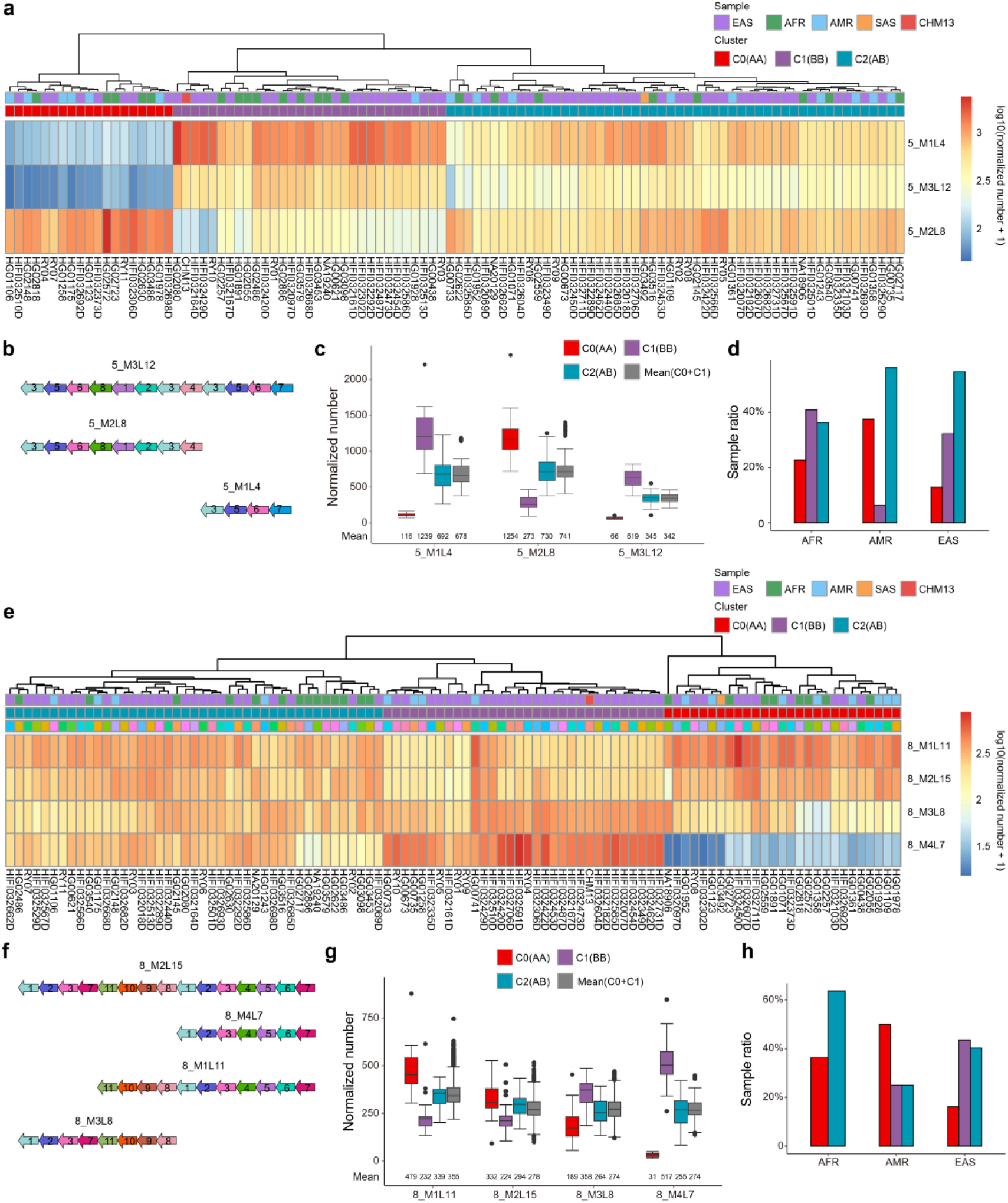
Centromere genotype from sample clustering based on HOR n-numbers in chromosome 5 and 8. **a.** The heatmap and sample hierarchical clustering of HOR n-numbers in chr5. **b.** Monomer patterns of 5_M3L12, 5_M2L8 and 5_M1L4. **c.** The box plot of HOR n-numbers in 5_C0 (AA), 5_C1 (BB) and 5_C2 (AB). For each HOR, the Mean(C0+C1) represents the pairwise mean n-numbers in 5_C0 and 5_C1. **d.** The proportion of samples in each of the AFR, AMR and EAS populations containing 5_C0, 5_C1 and 5_C2. **e.** The heatmap and sample hierarchical clustering of HOR n-numbers in chr8. **f.** Monomer patterns of 8_M2L15, 8_M1L11, 8_M3L8 and 8_M4L7. **g.** The box plot of HOR n-number among 8_C0 (AA), 8_C1 (BB) and 8_C2 (AB). For each HOR, the Mean(C0+C1) represents the pairwise mean n-numbers in 8_C0 and 8_C1. **h.** The proportion of samples in each of the AFR, AMR and EAS populations containing 8_C0, 8_C1 and 8_C2.

### Chromosome specific HOR landscapes

To develop our analysis beyond a population-level characterisation of HOR diversity, we next investigated the distribution of HOR loci between samples. We applied HiCAT-human-assembly to 109 assemblies and annotated the HOR patterns in each CASA. We then compared the distribution of HORs in CASAs between assemblies and found that the majority of chromosomes contained a discernible, and different, composition of HORs (hereafter ‘landscape’) and that 10 chromosomes seemingly contained two distinct landscapes (Fig. 4a, b and Supplementary figure S18, 19). For example, in chr11, the first of two landscapes is homogeneous, comprised entirely of one HOR (specifically, M1L5) with a repeating pattern of monomers of the form 1-2-3-4-5 (Fig. 4a). By contrast, the second landscape demonstrates the expansion of a different HOR, M2L1, within a larger set of M1L5 arrays; this arose by the tandem duplication of the first monomer in M1L5, which in this landscape has a repeating pattern of monomers of the form 1-1-2-3-4-5. This landscape of HORs has previously been detected in CHM1 ^8^ (Fig. 4a and Supplementary figure S20). We make similar observations for the CASAs of chr3, 6, 12, 14 and 20 (Supplementary figure S18a, c, f and S19a, e). The common feature for dual-landscape centromeres is that one landscape is homogeneous (dominated by one HOR) whereas the other is ‘locally nested’, showing the local expansion of a subunit within the primary HOR unit. We also found that the local expansion rates differ greatly between chromosomes, appearing relatively high for chr11 but lower for chr3 and 20 (Supplementary figure S21).

**Fig. 4.**
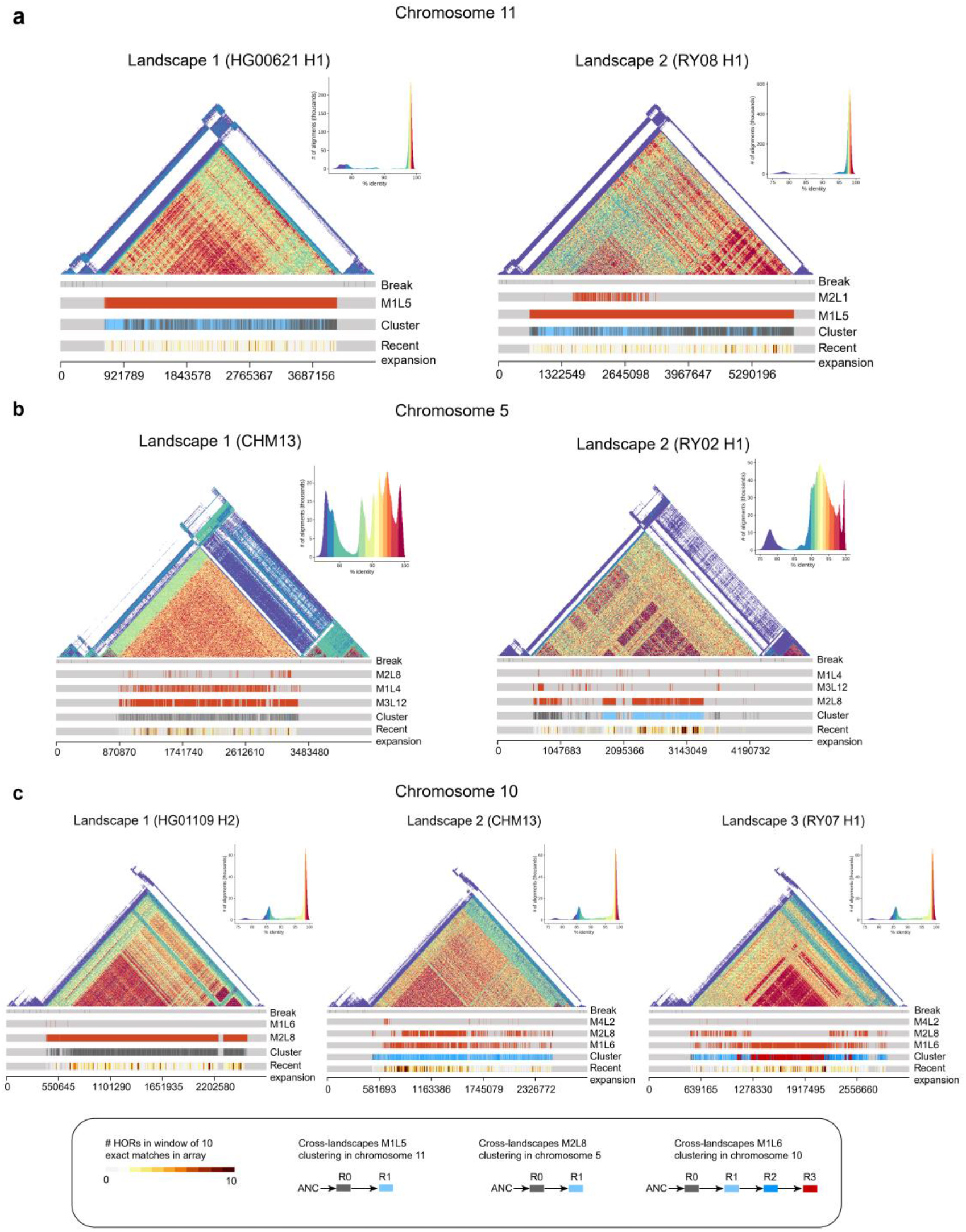
HOR landscapes on chromosome 11, 5 and 10. The HOR landscapes on chr11 represented by HG00621 haplotype 1 (H1) and RY08 H1 (**a**), chr5 represented by RY02 H1 and CHM13 (**b**), and chr10 represented by HG01109 H2, CHM13 and RY07 H1 (**c**). The triangle similarity heatmaps are generated by StainedGlass ^17^. The “Break” track shows the breakpoints in chromosome specific alpha satellite arrays (CASAs). The “Cluster” track represents the represents cross-landscape CASA HOR clustering results. The “Recent expansion” track represents HOR exact matching number within a sliding window of 10 HOR units (with 1 HOR slide). The other tracks record the position of different HORs. Ancestral HOR sequence (ANC) of each chromosome is reconstructed by monomers from other chromosome with the same suprachromosomal family ^10^.

In chr5, variation in HOR composition appeared even greater than the six aforementioned chromosomes (Fig. 4b). The primary HOR in landscape 1 (sample number n = 57) is M3L12 while that in landscape 2 (n = 52) is M2L8. Since M2L8 MP is a subunit of M3L12, we performed cross-landscape M2L8 clustering and found that cluster R0 in landscape 2 is shared with landscape 1 while cluster R1 is specifically enriched in landscape 2, constituting a new layer within it (Fig. 4b and Supplementary figure S22).

We found three types of landscape in the CASA of chr10 with landscape 1 primarily comprising HOR M2L8 (monomer pattern: 1-2-3-4-5-6-8-7) with a small number of M1L6 (monomer pattern: 1-2-3-4-5-6), landscape 2 showing the co-occurrence of M2L8 and M1L6 across the entire CASA, and landscape 3 similarly showing the co-occurrence of M2L8 and M1L6 although with M1L6 expanding within the middle (Fig. 4c). Cross-landscape M1L6 clustering shows that cluster R0 concentrates on landscape 1 and both ends of landscape 2 and 3. R1 and R2 are interlaced in landscape 2 and both sides of landscape 3. R3 specifically exists in middle of landscape 3 (Fig. 4c and Supplementary figure S23). We compared the consensus sequences of three clusters with the reconstructed ancestral sequence and found that R0 is the most similar to it, while R3 may have more recently expanded (Supplementary table S16). Based on this result, we proposed a model to illustrate the evolution of chr10 HORs. The ancestral landscape may be homogeneous with M2L8 (landscape 1) with a deletion or local duplication event within it having given rise to M1L6 which expanded by the process of ‘layer expansion’ ^7^ to form, initially, landscape 2, and then, after more extensive expansion, landscape 3.

We detected four types of landscapes of HOR in chr8 (Supplementary figure S18e). We found most samples (n = 62) represented by landscape 1, where M1L11 concentrated on both sides of CASA while M2L15 enriched in the middle. Landscape 2 (n = 37) represented by CHM13. Different from landscape 1, it has a recent expansion of M4L7 in the middle of CASA. Except for these two major landscapes, we also detected two minor landscapes: landscape 3, which was only found in 9 samples with M1L11 concentrated to the right of the CASA and M2L15 (with a locally nested M3L8) to the left; landscape 4 is similar to landscape 1 but the centre contains multiple copies of M3L8 and was only found in one sample (NA18906 H2).

Except for above chromosomes, other chromosomes contained only one HOR landscape among all samples, and they are grouped into two types (Supplementary figure S24-S26). The first one includes chr19, 22 and X and their CASAs are quite homogeneous dominated by a single HOR and similar among all samples. The second one contains a large number of locally nested HORs (LN-HOR), like chr1, 2, 9 and 15. In summary, these results demonstrate that while human centromere HOR arrays are diverse, they share structural resemblances in their composition and so form a number of ‘landscapes’.

### Different levels of locally nested HOR contributed to the landscape of centromere evolution

The above analyses suggest that LN-HOR play a recurring role in the evolution of centromeres. The CASA of chr1 primarily comprises 1_M2L6 (monomer pattern, MP: 1-2-5-6-4-3), with locally nested 1_M1L2 (a dimer, i.e., with repeating pattern 1-2) (Fig. 5a, Supplementary figure S25a and Supplementary table S17). We have found that the dimer expansion peak of 1_M2L6 is at the position of repeating four times with 12 monomers (1-2)×4-5-6-4-3 ^13^. We further analysed the monomer length distribution of all M2L6 units and found that there were different peak positions among all samples (Fig. 5b and Supplementary figure S27). The peaks in 68.6% samples are at the position of repeating three times with 10 monomers (1-2)×3-5-6-4-3, and the sample ratio with the peak at the position of 12 monomers ((1-2)×4-5-6-4-3) is higher in EAS (32.3%) than that in AFR (18.2%, *p*-value is 0.005, one-sided binomial test) or AMR (18.8%, *p*-value is 0.008) (Supplementary figure S27g).

**Fig. 5.**
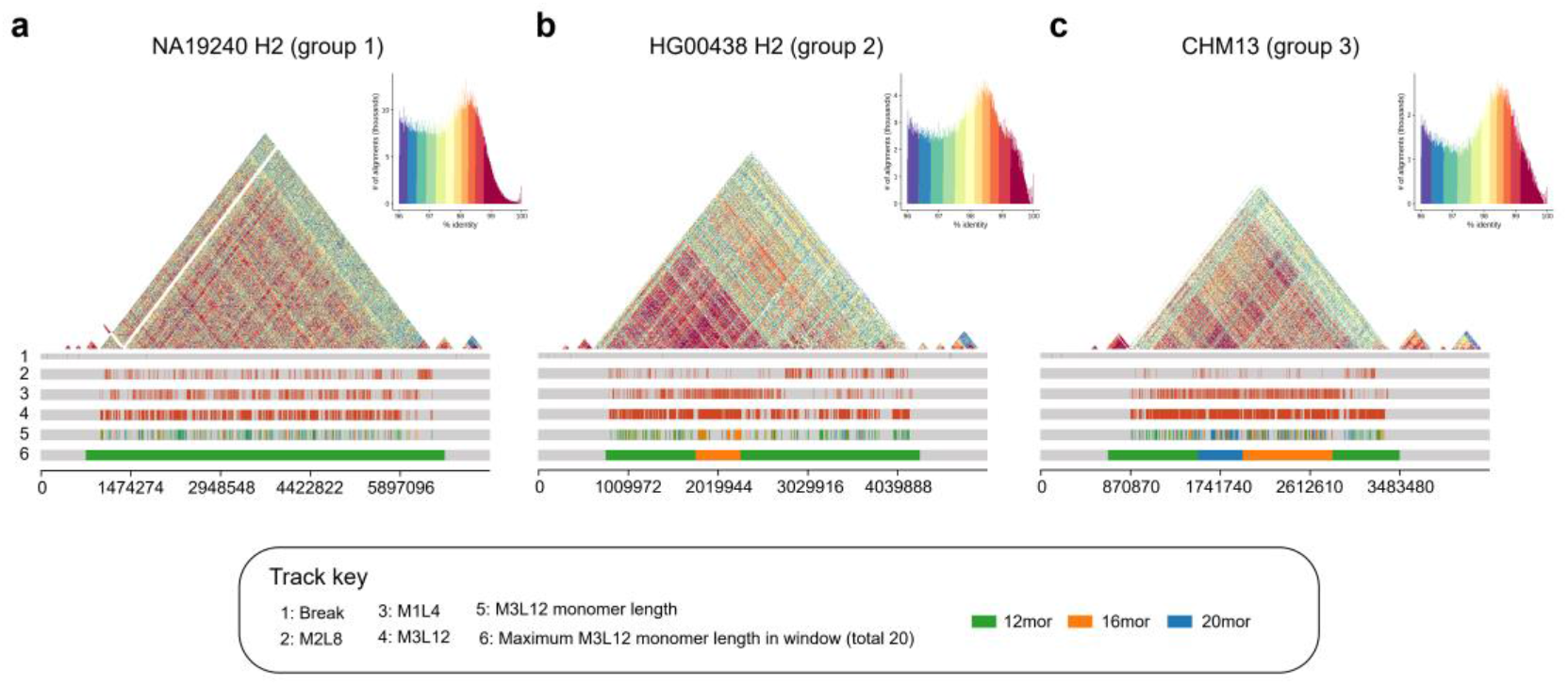
Locally nested HOR variation in chromosome 5 centromere arrays. Chr5 CASA landscapes represented by NA19240 H2 (**a**), HG00438 H2 (**b**) and CHM13 (**c**). The triangle similarity heatmaps are generated by StainedGlass ^17^ and exclude those sequences with identity lower than 96%. The first track shows the breakpoints in CASAs. The second to fourth tracks show the position of 5_M2L8, 5_M1L4 and 5_M3L12. The fifth track shows the position of 12 monomers (12mor, green), 16mor (orange) and 20mor (blue) of M3L12 units. In the sixth track, the whole CASA is split into 20 windows with each window showing the maximum value of the total number of 12mor, 16mor and 20mors.

Our previous study on CHM13 centromere HORs found that the 5_M3L12 units in chr5 centromere have three frequent MPs with 12 monomers (12mor, 3-5-6-8-1-2-3-4-(3-5-6-7)×1), 16mor (3-5-6-8-1-2-3-4-(3-5-6-7)×2) and 20mor (3-5-6-8-1-2-3-4-(3-5-6-7)×3) ^13^. We wonder whether these three MPs have different frequency among samples with different patterns. The CASA of CHM13 chr5 belongs to landscape 1, where 5_M3L12 exist with M1L4 expansion in the whole array. Study on landscape 1 shows that the ratio distributions of these three MPs from M3L12 can be clustered into three groups. In group 1 (27 samples), the ratio of 12mor is high and that of 16mor and 20mor is low, while in group 2 (18 samples), the ratio of 16mor is high relative to group 1. In group 3 (12 samples) including CHM13, all three MPs have a high frequency (Supplementary figure S28 and Supplementary table S18). Taking NA19240 H2 (group 1), HG00438 H2 (group 2) and CHM13 (group 3) as examples, the entire CASA of NA19240 H2 is represented by a 12mor MP while the middle region of the HG00438 H2 CASA is enriched with 16mor (Fig. 5a, b). Different from above two samples, a region dominated by a 20mor appears to the left side of the 16mor region in the CHM13 CASA (Fig. 5c). These results suggest that different levels of LN-HORs occur in human centromeres, and that with subsequent mutations these may ultimately contribute to HOR landscape differentiation.

## Discussion

We investigated the population diversity of human centromere sequences based on both HiFi reads and haplotype-resolved assemblies of hundreds of samples from two pangenome sequencing consortiums (HPRC and CPC). We reported considerable diversity in centromere HOR arrays size for different samples, with CHM13 representing only one of many possible human HOR patterns. In addition, we found 33 HORs showed variable numbers in all samples and 23 of them show significantly different distributions among three populations. We detected three centromere genotypes with imbalance population frequencies on chr5, 8 and 17. Moreover, a comparative analysis of CASAs across assemblies revealed that although human centromere HOR array structures are diverse, they nevertheless tend to resolve into a relatively small number of landscapes, with LN-HORs playing an important role in their diversification.

In a previous analysis of the b/n boxes (CENP-B-binding site and pJα protein-binding site) within the HOR units of a single genome, a cyclical model was proposed to explain how HOR units vary among chromosomes, structured around three states with short, moderately longer and substantially longer HOR units, respectively ^18,19^. Under this model, the rate of tandem expansion within longer HOR units is higher than that of shorter HOR units, but that shorter HOR units have a more stable b/n box structure and so can better resist invasion by pericentromeric heterochromatin. This degree of competition between the rate of tandem expansion (which determines HOR length) and its capacity to resist invasion (which determines b/n box completeness) influences which HOR expands, and how greatly.

However, how do centromere HOR arrays evolve in the broader human population? In this study, based on the analysis of hundreds of human genomes, we reported considerable diversity of HOR landscapes for each chromosome which may be explained by a similarly cyclical model centred on the transition of HOR landscapes between homogeneous and locally nested states (Fig. 6). In a homogeneous landscape, the centromere is mainly dominated by a single HOR, like that observed on chr19, 22 and X. The recurrent expansion of these HOR subunits eventually ‘breaks’ the homogeneous landscape (through replication error such as a monomer duplication), forming locally nested units which have the advantage of a higher expansion rate (because by disproportionately occurring in the middle of existing HORs, they are protected from invasion by pericentromeric heterochromatin). These newly expanded subunits may increase the density of the b box, similar to our observations on chr11 landscape 2 (Supplementary figure S29a), obtaining a competitive advantage relative to the homogeneous state (we call this the ‘seeding stage’). Following centromere drive ^20^, LN-HORs rapidly expand and so spread throughout the population (‘expansion stage’). These two stages may be typified by chr3, 11, 12, 14 and 20, each of which has dual-landscape centromeres, one being highly homogeneous and the other with varying levels of local nesting. LN-HOR expansion results in a state with high LN-HOR content in array and high LN centromeres ratio in population, represented by chr1, 2, 9 and 15 (‘locally nested state’). Once locally nested state is formed, the entire array is composed by original HORs, the LN subunits and their combinations with different levels. These HOR units containing different monomer length and b box content may achieve different advantages for competition, leading to rapid and pronounced changes in HOR landscapes, like chr5 and chr8 (‘differentiation stage’). Ultimately, one type of HOR unit may ‘win’ in the competition and gradually homogenize the landscape (‘homogenization stage’), whereupon the cycle repeats.

**Fig. 6.**
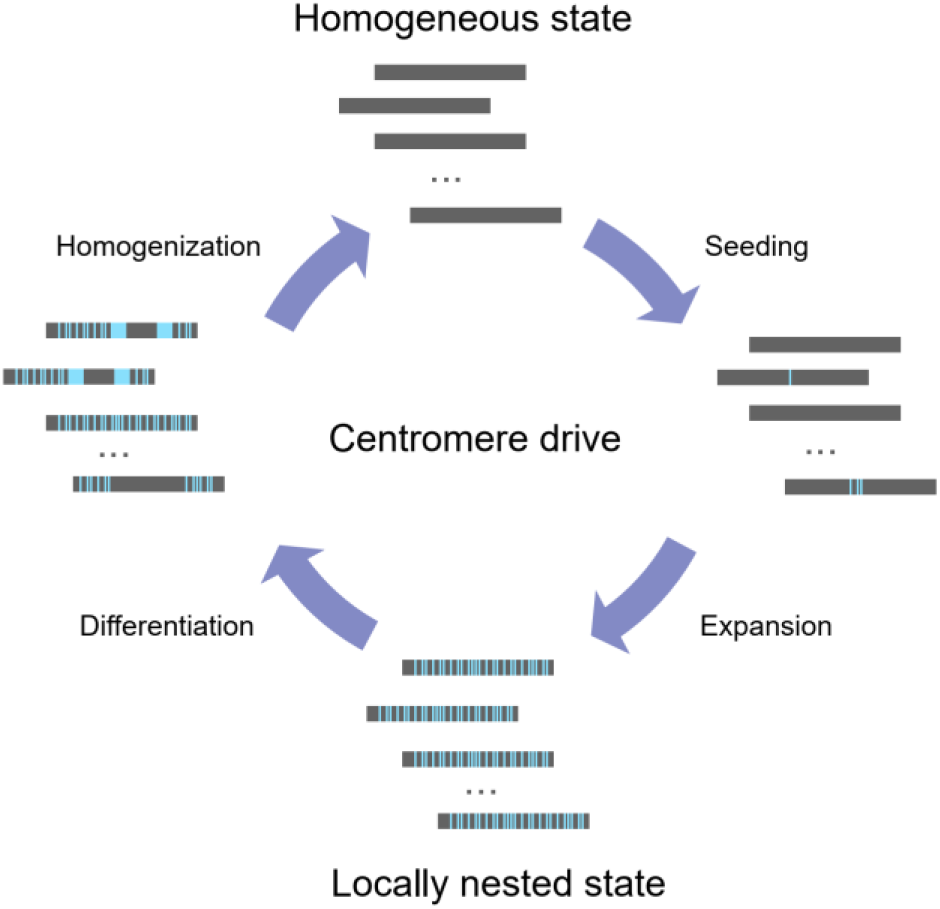
Cyclical model of HOR landscape evolution in human population. In the homogeneous state, all HOR arrays are homogeneous with only one HOR large expansion. In seeding stage, variation occurs in original HORs forming the LN-HOR units (seeding stage). In expansion stage, under the centromere drive ^20^, LN-HOR units expand rapidly and centromeres with large number of LN-HOR units spread in the population forming locally nested state. In differentiation stage, HORs with different monomer length and b box content compete in the array leading to different or even dramatical changes in HOR landscapes. Finally, one type of HOR units may “win” the competition and gradually back to homogeneous state (homogenization stage).

Three HOR landscapes in chr10 may record this cycle (Fig. 4c). Landscape 1 (n = 6 samples) consists entirely of 10_M2L8, representing the homogeneous state. In landscape 2 (n = 59), 10_M2L8 and 10_M1L6 (subunit of 10_M2L8) coexists across the entire CASA, representing a locally nested state. For landscape 3 (n = 44), a significant expansion of 10_M1L6 is observed in the middle of array with co-occurrence of M1L6 and M2L8 on either side, which may represent a period of transition from a locally nested to a homogeneous state. We observed a high density of b boxes in the newly expanded 10_M1L6 on landscape 3 which may also provide support for the presumed selective advantage of this state (Supplementary figure S29b).

In summary, using a large number of samples with high-coverage HiFi reads data and high-quality haplotype-resolved assemblies, we demonstrated the diversity of human centromere HOR patterns and explored how they evolved. In the future, the release of more human genomes and the accumulation of related epigenetic data has the potential to refine this model, and so may provide further insights into our understanding of the mechanisms of centromere evolution.

## Methods

### HiFi sequencing data and genome assembly

We obtained HiFi reads for 43 samples released by the HPRC ^12^, 58 samples by the CPC ^11^, plus CHM13 ^2^. All HiFi reads data were first converted into fasta format using seqtk (v1.3-r106, https://github.com/lh3/seqtk). Samtools (v1.12) was used to calculate read length for each sample ^21^. We also obtained 22 contig-level haplotype-resolved assemblies (11 samples) from CPC and 86 assemblies (representing 43 individuals) from HPRC, and CHM13 genome ^2,11,12^. Accession numbers and sample metadata for both HiFi reads and assemblies are given in Supplementary Table S1. For each contig-level assembly, we used RagTag (v2.1.0) ^22^ compared with CHM13 genome to generate the chromosome-level assembly. For each HPRC assembly, haplotypes 1 and 2 (H1 and H2) represent the paternal and maternal assembly, respectively.

### Building chromosome-specific alpha satellite arrays

For each chromosome of each assembly, alpha satellite units were identified by using Lastz (v1.04.15) ^14^ to map the alpha satellite template sequence. Alpha satellite units mapped < 5kb from each other were merged to produce a set of alpha satellite regions and the regions with total length < 10kb were discarded. Finally, for each chromosome, we concatenated its set of alpha satellite regions, producing its CASA.

### Alpha satellite reads classifier training

HiCAT-human-reads contains a two-step classifier based on principal component analysis (PCA) and support vector machine (SVM) for extracting chromosome-specific alpha satellite reads (ASRs). The first step was detecting ASRs and the second step was assigning ASRs to their chromosome of origin. Firstly, a training dataset was simulated using the CHM13 genome with HG002 chrY (CHM13-HG002 chrY) ^2,23^. To simulate ASRs, CASAs from CHM13-HG002 chrY were randomly broken into reads with length of 10-25 kb, of which we reversed half of the set to represent the “-” strand. We repeated the simulation 10 times. To ensure a balanced dataset, the number of ASRs simulated from each chromosome were equal. We simulated the same number of negative (non-ASR) samples for training by randomly sampling 10-25 kb sub-sequences from CHM13-HG002 chrY with the exception of alpha satellite regions. Simulated ASRs were labeled as “Alpha” and non-ASRs as “Non-Alpha” for the first step, with the former labeled according to their corresponding chromosomes for the second step. Secondly, we calculated the frequency of all 5-mers to construct a 5-mer frequency matrix *M_n_*_×_*_m_* for all simulated reads, where *n* denotes read count and *m* = 512 denotes all 5-mers on each of the two (“+/-”) strands (1024 / 2 = 512). We normalized the 5-mer frequency matrix by the corresponding read length, then used PCA to reduce the dimension of the normalized 5-mer frequency matrix while ensuring that the amount of variance it explains exceeds 95%. Finally, we adopted a SVM to classify reads.

### Running HiCAT and monomer template selection

We ran HiCAT (v1.1) ^13^ on each CASA from CHM13-HG002 chrY to annotate its monomers and HORs. For each CASA, HiCAT annotates its constituent monomers using a community detection approach ^24^. In brief, for a monomer community with more than one monomer, HiCAT calculates pairwise edit distances between monomers, then selects the monomer sequence with the lowest edit distance to all other monomers in the community as its output (that is, the template sequence from which HORs are built).

### HOR annotation

In HiCAT-human annotation, we first used the StringDecomposer algorithm ^15^ to decompose ASRs and CASAs into block sequences. Then, for each chromosome, we calculated the sequence identity between each block and the monomer template (obtained by HiCAT annotation of CHM13-HG002 chrY, described above). The sequence identity was defined as:

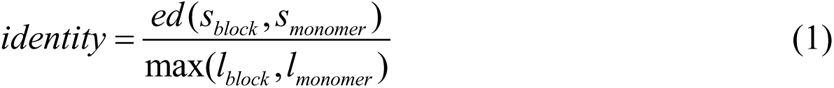

where *ed*(*s_block_*, *s_monomer_*) is the edit distance between the block and monomer template sequence. *l_block_* is the sequence length of the block and *l_monomer_* is the sequence length of monomer template. We labelled the block with the largest identity monomer ID to transform block sequences into monomer sequences. We classified blocks whose highest identity was lower than 90% as ‘unknown sequence’; this may represent transposable elements or other sequence interjected into the HOR. Since locally nested HORs occur in human centromeres, HiCAT-human adopted the HTRM method which we previous developed ^13^ to recursively detect and compress local tandem repeats in the monomer sequences to obtained HORs in each chromosome. This has the effect of ensuring that HORs with shifted or reversed units, such as those with monomer patterns 1-2-3-4, 4-1-2-3, 3-4-1-2, 2-3-4-1 and 4-3-2-1, will be grouped together as the same HOR. For presentation purposes, HORs were sorted on the basis of their total number of repeats and then named following the convention “R + rank + L + unit monomer length”.

### Multi-sample HOR aggregation

To compare cross-sample annotations, we aggregated the annotation results for both HiFi reads and assemblies. For HiFi reads, HORs with shifted and reversed units were grouped together with the same HOR pattern (as described above). HOR patterns were sorted by the total repeat number across all samples and named following the convention “M + rank + L + unit monomer length”. For assembly, HOR pattern names are matched with HiFi reads.

### Quantification of HiFi reads HORs among all samples

The estimated HOR array size *s_i_*_,*k*_ of chromosome *i* in sample *k* is defined as:

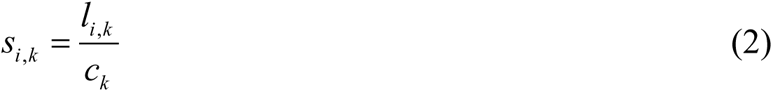

where *l_i_*_,*k*_ is total length of HiFi reads with HOR of chromosome *i* in sample *k* and *c_k_* is the sequencing coverage of sample *k*.

The n-number *n_j_* _,*k*_ of HOR *j* and sample *k* is calculated as following:

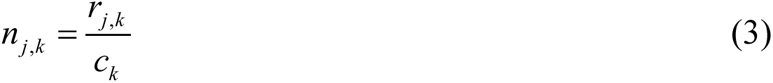

where *r_j_* _,*k*_ represents the number of HOR *j* in sample *k* output by HiCAT-human-reads. We excluded the rare HORs which meet the following condition:

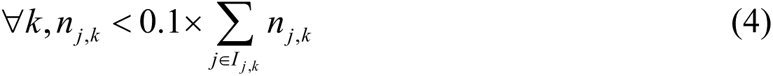

*I _j_* _,*k*_ represents all HORs on HOR *j* corresponding chromosome of sample *k*. For the remaining HORs, the mean-fold change *mf j* _,*k*_ of HOR *j* in sample *k* is calculated as following:

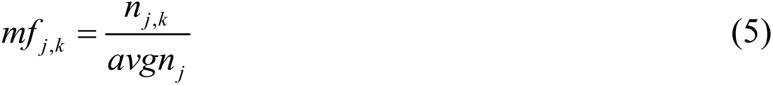

where *avgn_j_* is the mean number of HOR *j* among samples. Then, we calculated the standard deviation*std _j_* of the mean-fold change of HOR *j* among samples. The v-HOR is defined as HOR *j* with *std _j_* larger than 0.5. We obtained 33 v-HORs.

### Cross-landscape HOR clustering

To compare the HOR among different landscapes, we performed a cross-landscape HOR clustering analysis (Supplementary figure S30). For different landscapes, the largest monomer pattern that their primary HORs shared are extracted. For example, in chr5, the primary HORs are M3L12 in landscape 1 and M2L8 in landscape 2. Since M2L8 is subunit of M3L12, we selected M2L8 as target HOR in chr5 cross-landscape HOR clustering. We extracted all DNA sequences of target HOR units in different CASAs and performed multiple alignment for these sequences using Kalign (v3.3.5) ^25^. Then, the most common base at every position of the alignment files was identified to build the target HOR consensus sequence. All extracted HOR DNA sequences were pairwise aligned with the consensus sequence (needle, EMBOSS v6.6.0)^26^ and reformatted into 0-1 vectors, where 0 indicates that the HOR shares the same base with the consensus sequence at that position and 1 indicates there is a difference. The 0-1 vectors of all HOR units were clustered based on k-means. For each target HOR, we chose the smallest k that can show the difference among landscapes.

### Ancestral HOR sequence reconstruction

To explore the evolution of HOR clusters among different landscapes, we performed ancestral HOR sequence (ANC) reconstruction and compared the HOR clusters consensus sequences with ANC. For each target chromosome, we used monomers from another chromosome in the same suprachromosomal family (SF) as outgroup to reconstruct ANC ^10^. For chr5 and 10 (SF1), we used monomers from chr19 since its primary HOR has the same unit monomer length as the ancestor and for chr11 (SF3), we used chr17. For each outgroup chromosome, we generate the consensus sequence of each monomer of the primary HOR and built a monomer phylogenetic tree using IQ-TREE (v2.2.5) ^27^. The monomers were divided into groups and the group number was based on the SF ancestral HORs. We calculated the consensus sequences of monomers from each group as ancestral monomers. Then, for each target chromosome, we generated the consensus sequences of each monomer in the target HOR. We used these sequences with the ancestral monomers to build a phylogenetic tree using IQ-TREE, obtaining a correspondence between ancestral monomers and target monomers. We then derived the ANC DNA sequence based on ancestral monomers and target HOR. Finally, we used Clustal Omega ^28,29^ to perform a multiple sequence alignment of the ANC with the consensus sequences of each HOR cluster, inferring their relationship from the identity matrix.

## Data availability

This study used published data for analysis. Accession numbers and sample metadata for both HiFi reads and assemblies are given in Supplementary Table S1. The assembly and HiFi sequencing data of CHM13 human cell line can be accessed from GitHub at https://github.com/marbl/CHM13#downloads. The assembly and HiFi sequencing data of HPRC samples can be accessed at https://s3-us-west-2.amazonaws.com/human-pangenomics/index.html. The assembly of CPC samples can be accessed at https://ngdc.cncb.ac.cn/bioproject/browse/PRJCA011422. The HiFi sequencing data of CPC samples is obtained from Fudan University. The HOR annotation results and the HOR landscapes of all samples are deposited at https://figshare.com/articles/dataset/HiFi_reads_and_assemblies_HOR_annotation/25067558.

## Code availability

HiCAT-human code has been deposited at https://github.com/xjtu-omics/HiCAT-human and zenodo DOI: https://zenodo.org/doi/10.5281/zenodo.10570850. The cross-landscape HOR clustering analysis scripts have been deposited in at https://github.com/xjtu-omics/Cross_landscape_HOR_clustering and zenodo DOI: https://zenodo.org/doi/10.5281/zenodo.10570634.

## Acknowledgements

We thank Yang Gao and Shuhua Xu for data sharing, and Jing Hai and Huanhuan Zhao for administrative and technical support. This study was supported by the National Natural Science Foundation of China (32125009; 62172325; 32070663; 32200510); the National Key R&D Program of China (2022YFC3400300); the Natural Science Foundation of Shaanxi Province (2024JC-JCQN-28).

## Author contributions

KY and XY conceived the study. SG, YZ, XY and BW and analysed the data. SG and YZ developed the program. SG, XY, SB and YZ wrote the manuscript. SG, XY and YZ completed figures of manuscript. All authors read and approved the final manuscript.

## References

1. Barra, V. & Fachinetti, D. The dark side of centromeres: types, causes and consequences of structural abnormalities implicating centromeric DNA. Nat Commun 9, 4340 (2018).

2. Nurk, S. et al. The complete sequence of a human genome. Science 376, 44–53 (2022).

3. McNulty, S.M. & Sullivan, B.A. Alpha satellite DNA biology: finding function in the recesses of the genome. Chromosome Res 26, 115–138 (2018).

4. Bzikadze, A.V. & Pevzner, P.A. Automated assembly of centromeres from ultra-long error-prone reads. Nat Biotechnol 38, 1309–1316 (2020).

5. Jain, C., Rhie, A., Hansen, N.F., Koren, S. & Phillippy, A.M. Long-read mapping to repetitive reference sequences using Winnowmap2. Nat Methods 19, 705–710 (2022).

6. Wenger, A.M. et al. Accurate circular consensus long-read sequencing improves variant detection and assembly of a human genome. Nat Biotechnol 37, 1155–1162 (2019).

7. Altemose, N. et al. Complete genomic and epigenetic maps of human centromeres. Science 376, eabl4178 (2022).

8. Logsdon, G.A. et al. The variation and evolution of complete human centromeres. bioRxiv (2023).

9. Suzuki, Y., Myers, E.W. & Morishita, S. Rapid and ongoing evolution of repetitive sequence structures in human centromeres. Sci Adv 6(2020).

10. Romanova, L.Y. et al. Evidence for selection in evolution of alpha satellite DNA: the central role of CENP-B/pJ alpha binding region. J Mol Biol 261, 334–40 (1996).

11. Gao, Y. et al. A pangenome reference of 36 Chinese populations. Nature 619, 112–121 (2023).

12. Liao, W.W. et al. A draft human pangenome reference. Nature 617, 312–324 (2023).

13. Gao, S. et al. HiCAT: a tool for automatic annotation of centromere structure. Genome Biol 24, 58 (2023).

14. Harris, R.S. Improved pairwise alignment of genomic DNA, (The Pennsylvania State University, 2007).

15. Dvorkina, T., Bzikadze, A.V. & Pevzner, P.A. The string decomposition problem and its applications to centromere analysis and assembly. Bioinformatics 36, i93–i101 (2020).

16. Aldrup-MacDonald, M.E., Kuo, M.E., Sullivan, L.L., Chew, K. & Sullivan, B.A. Genomic variation within alpha satellite DNA influences centromere location on human chromosomes with metastable epialleles. Genome Res 26, 1301–1311 (2016).

17. Vollger, M.R., Kerpedjiev, P., Phillippy, A.M. & Eichler, E.E. StainedGlass: interactive visualization of massive tandem repeat structures with identity heatmaps. Bioinformatics 38, 2049–2051 (2022).

18. Rice, W.R. A Game of Thrones at Human Centromeres II. A new molecular/evolutionary model. bioRxiv, 731471 (2019).

19. Rice, W.R. A Game of Thrones at Human Centromeres I. Multifarious structure necessitates a new molecular/evolutionary model. bioRxiv, 731430 (2020).

20. Talbert, P.B. & Henikoff, S. What makes a centromere? Exp Cell Res 389, 111895 (2020).

21. Li, H. et al. The Sequence Alignment/Map format and SAMtools. Bioinformatics 25, 2078–9 (2009).

22. Alonge, M. et al. Automated assembly scaffolding using RagTag elevates a new tomato system for high-throughput genome editing. Genome Biol 23, 258 (2022).

23. Rhie, A. et al. The complete sequence of a human Y chromosome. Nature 621, 344–354 (2023).

24. Traag, V.A., Waltman, L. & van Eck, N.J. From Louvain to Leiden: guaranteeing well-connected communities. Sci Rep 9, 5233 (2019).

25. Lassmann, T. Kalign 3: multiple sequence alignment of large data sets. Bioinformatics 36, 1928–9 (2019).

26. Madeira, F. et al. The EMBL-EBI search and sequence analysis tools APIs in 2019. Nucleic Acids Res 47, W636–W641 (2019).

27. Minh, B.Q. et al. IQ-TREE 2: New Models and Efficient Methods for Phylogenetic Inference in the Genomic Era. Mol Biol Evol 37, 1530–1534 (2020).

28. Sievers, F. & Higgins, D.G. The Clustal Omega Multiple Alignment Package. Methods Mol Biol 2231, 3–16 (2021).

29. Madeira, F. et al. Search and sequence analysis tools services from EMBL-EBI in 2022. Nucleic Acids Res 50, W276–W279 (2022).

