## Supplementary information, figures and tables for "Centromere landscapes resolved from hundreds of human genomes": Supplementary information.pdf

**Supplementary note.**

**Simulation test for reads classification**

To examine the performance of the HiCAT-human-reads workflow, we simulated 30X HiFi sequencing data of CHM13 genome as basic test and the chromosome Y was from HG002 genome. The simulated sequencing error rate was 0.1% and read length ranged from 10 to 25Kb [1]. We repeated the simulation five times. We first evaluated the classification accuracy of all simulated alpha satellite reads (ASRs). We found that the average recall was 87.8% (Fig. S1a, Supplementary table S2). We found that most of false classified ASRs were classified into non-alpha regions (Fig. S1b). We further investigated the location of false classified ASRs and found they were concentrated on the flanking of active HOR regions (Fig. S1c). Chromosomes 13, 14, 15 and 21 each had a recall lower than 75%, which we found was driven by the false classification of ASRs in chromosomes 13, 14 and 15 as chromosome 21 due to segmental duplications (Fig. S1d). We summarized the HOR annotation of false classified ASRs and found that the average HOR coverage ratio (percent of HOR bases covered by a read) was only 4.3% and that 75.83% of the falsely classified ASRs did not have any HORs, indicating these reads have less impact on further HOR analysis (Supplementary table S3). Then, we examined the classification recall of simulated reads from active HOR regions defined in previous studies [2], and found the average recall was 99.7% (Fig. S1e, Supplementary table S2). We found that most falsely classified reads in active HOR regions were found on chromosomes 1, 13, 14 and 22 due to the fact that chromosomes 1/5/19, 13/21 and 14/22 each contained a shared set of HORs [2], and located in their marginal areas indicating that the marginal areas of active HOR regions were more similar inter chromosomes (Fig. S1f).

**Supplementary figures.**

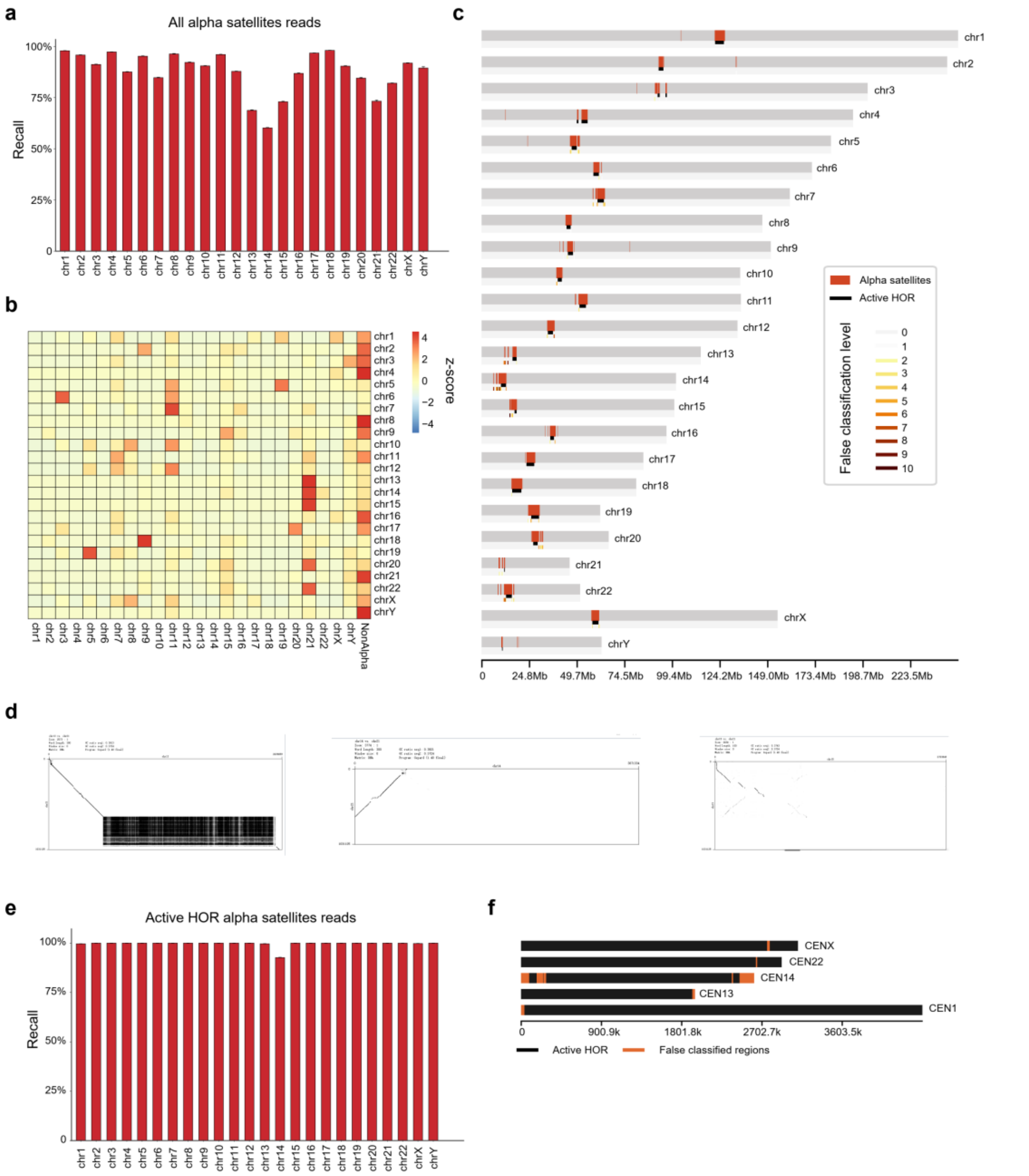


**Supplementary figure S1. Reads classification evaluation on simulated HiFi sequencing data. a.** Recall of reads classification in all alpha satellite regions for each chromosome. **b.** Heatmap of false classified reads in all alpha satellite regions and normalized based on z-score. **c.** The location and level of false classified reads in all alpha satellite regions. The window size is 500Kb. **d.** Dot plots between CHM13 chromosome 21 CASA with chromosome 13, chromosome 14 and chromosome 15. **e.**

Recall of reads classification in active HOR regions for each chromosome. **f.** The location of false classified reads in active HOR regions. CEN means active HOR regions in each chromosome centromere.

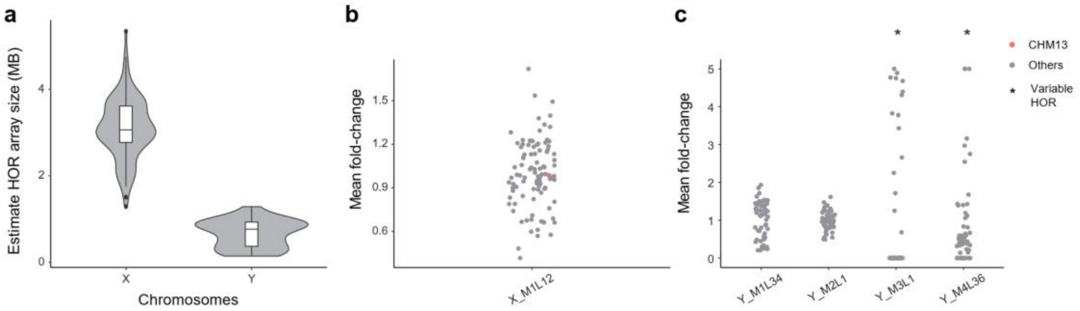


**Supplementary figure S2. HOR quantification on chromosome X and chromosome Y. a.** The estimate HOR array size based on total HOR read length and sequencing coverage. **b-c.** The variation of HOR mean fold-change on chrX (**b**) and chrY (**c**) among all samples. CHM13 is represented by red and other samples are grey. v-HORs are marked by stars.

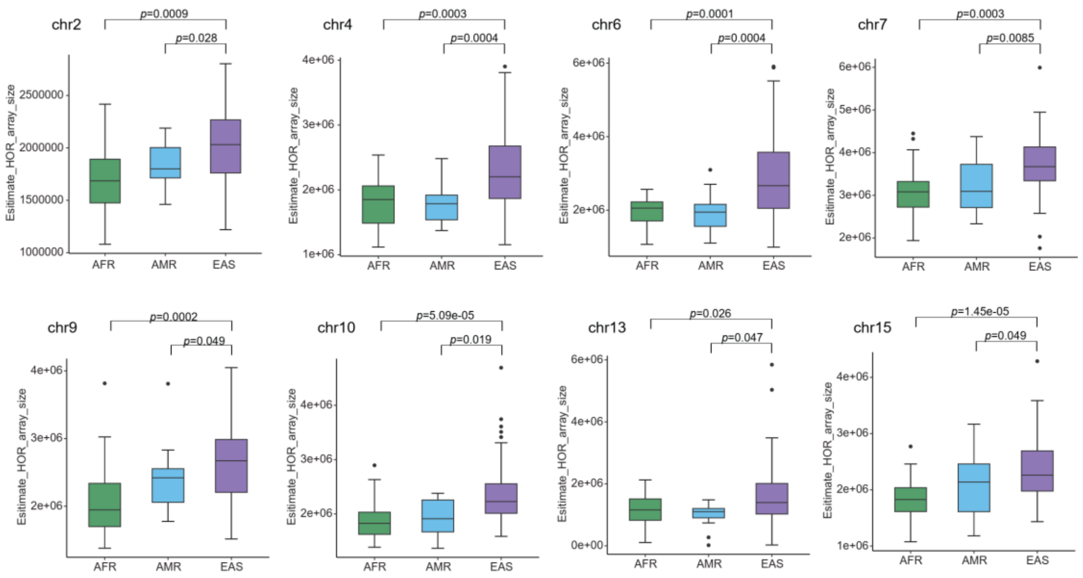


**Supplementary figure S3. Chromosome HOR array size in AFR, AMR and EAS significantly larger in EAS than other populations.** HOR array size is significantly larger in EAS samples than that in both AFR samples and AMR samples on chr2, 4, 6, 7, 9, 10, 13, 15. *p*-value is calculated by one-sided Wilcoxon rank sum test.

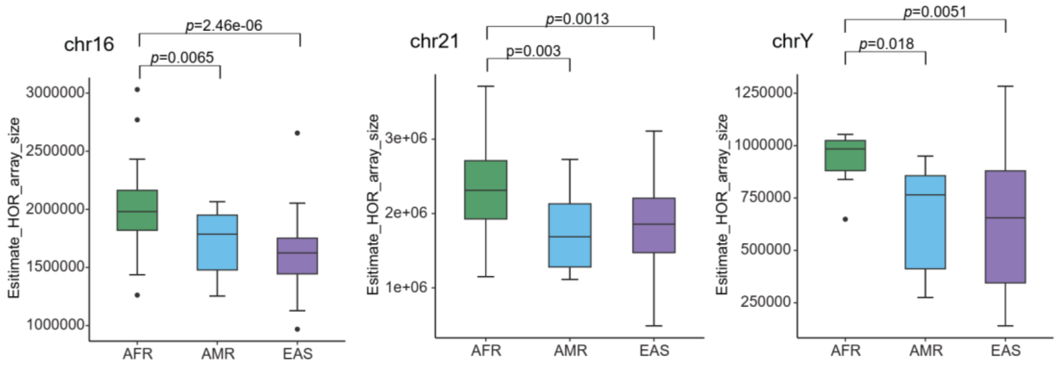


**Supplementary figure S4. Chromosome HOR array size in AFR, AMR and EAS significantly larger in AFR than other populations.** HOR array size is significantly larger in AFR samples than that in both EAS samples and AMR samples on chr16, 21 Y. *p*-value is calculated by one-sided Wilcoxon rank sum test.

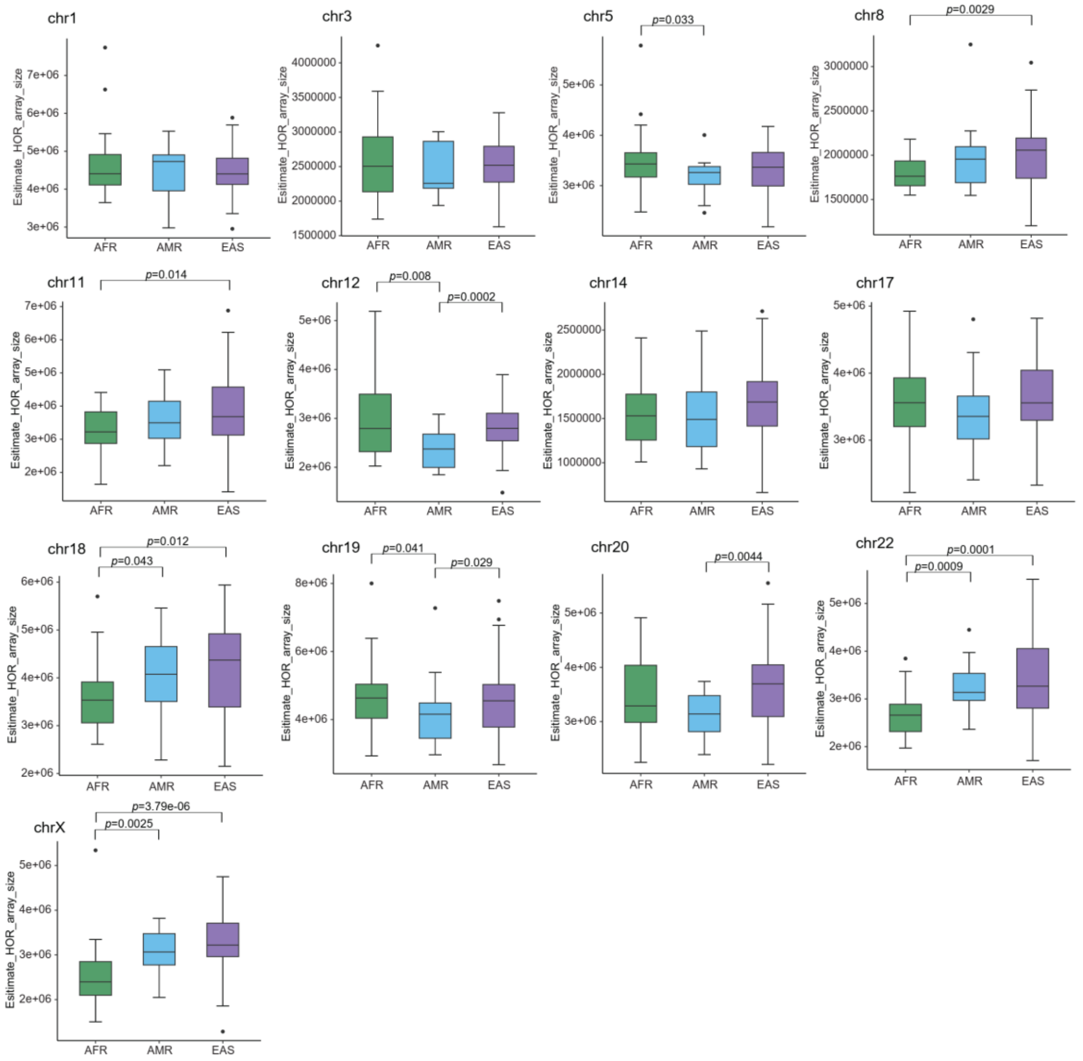


**Supplementary figure S5. Chromosome HOR array size in AFR, AMR and EAS on other chromosomes.** *p*-value is calculated by one-sided Wilcoxon rank sum test.

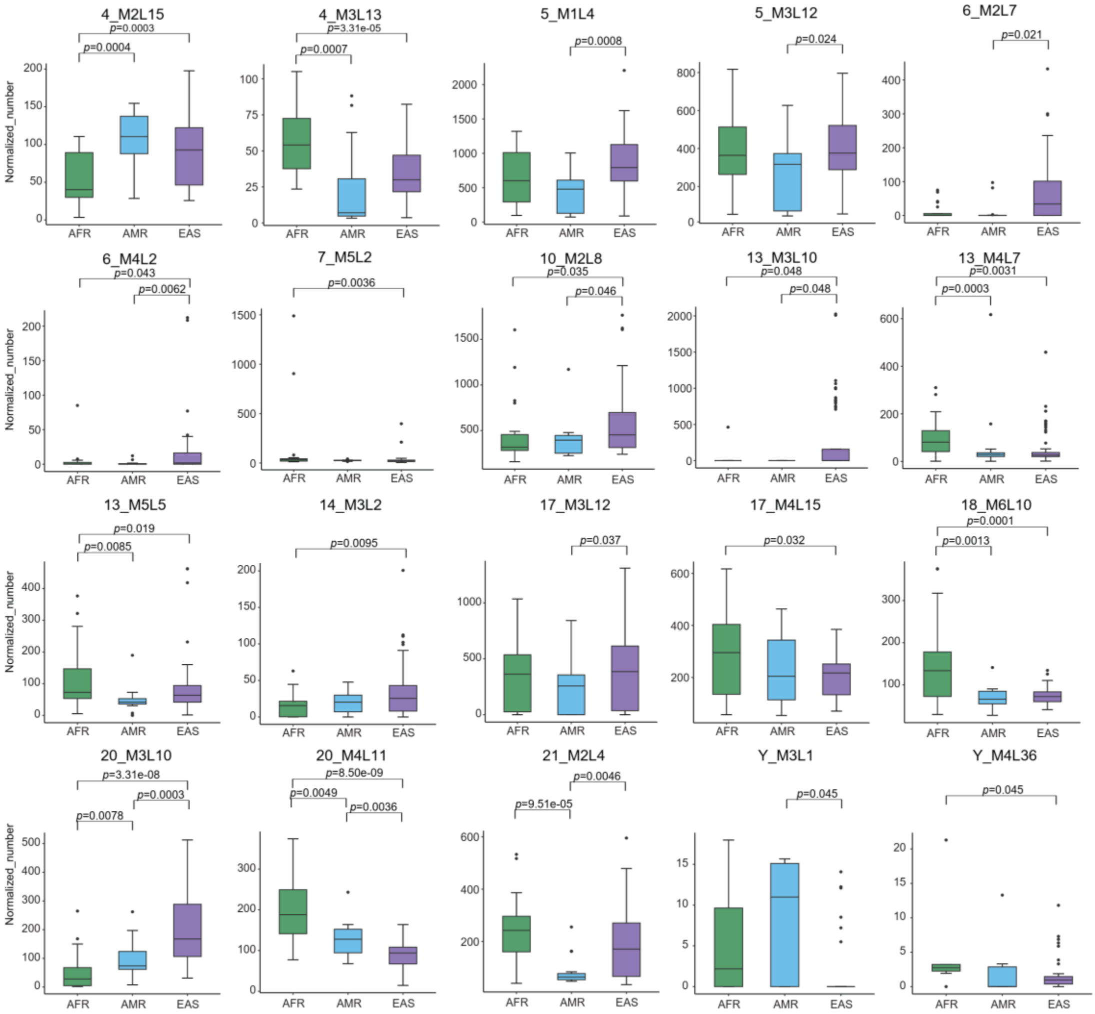


**Supplementary figure S6. v-HORs showing significant variance among populations.** *p*-value is calculated by two-sided Wilcoxon rank sum test.

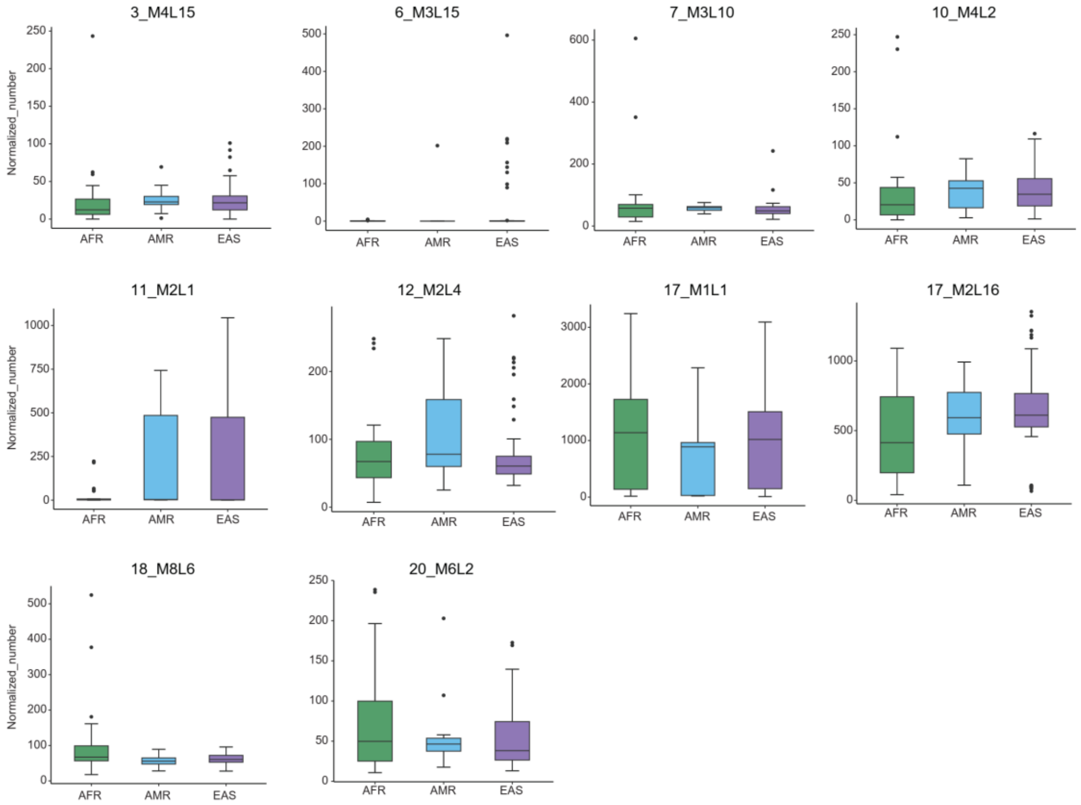


**Supplementary figure S7. v-HORs not showing significant variance among populations.** *p*-value is calculated by two-sided Wilcoxon rank sum test.

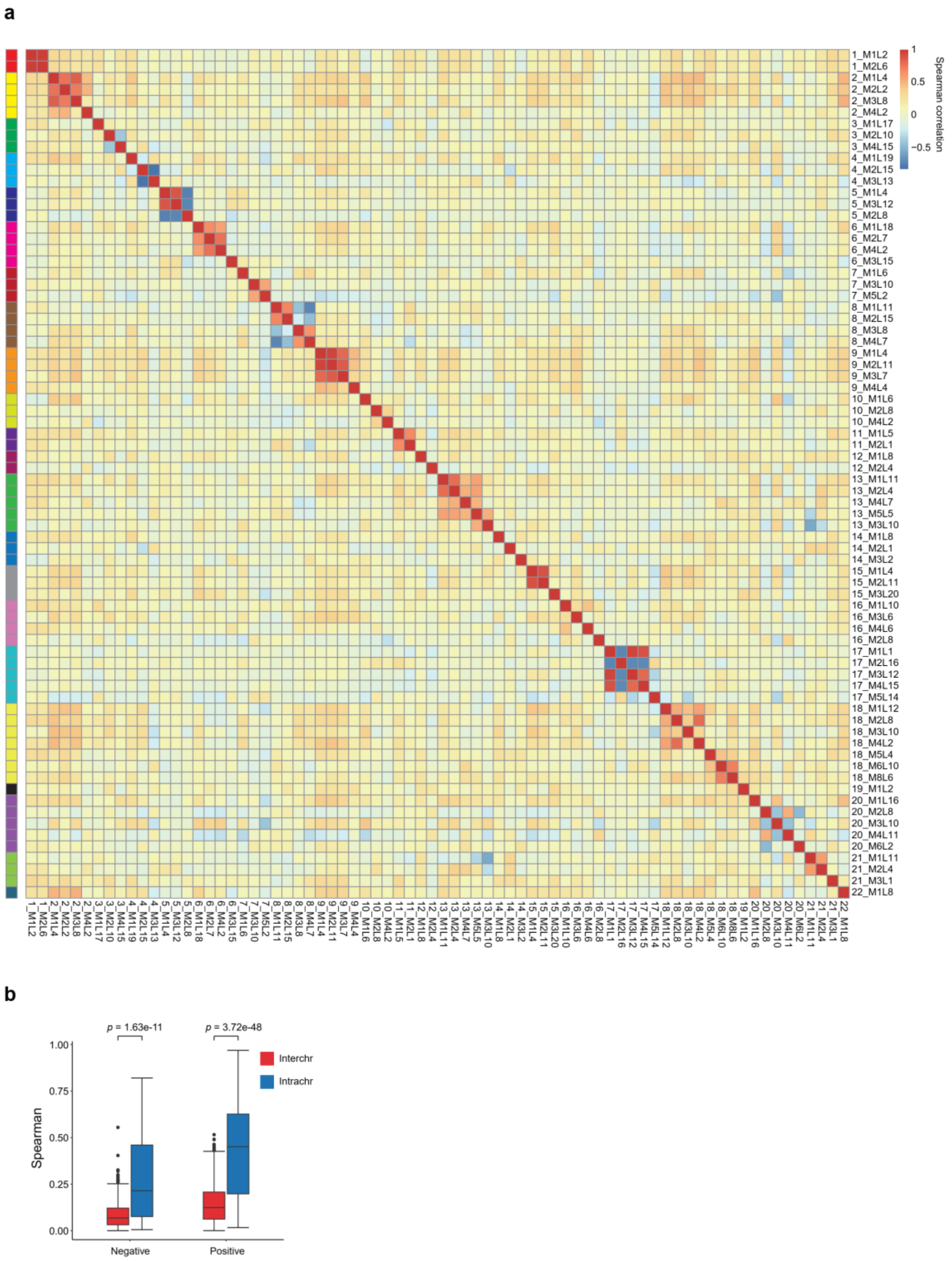


**Supplementary figure S8. Spearman correlation between all HORs. a.** The heatmap shows the correlation of HOR n-numbers. The different colour in left bar indicates the HOR from different chromosomes. **b.** Absolute value of n-number spearman correlation coefficient of inter and intra chromosome HORs. *p*-value is calculated by one-sided Wilcoxon rank sum test.

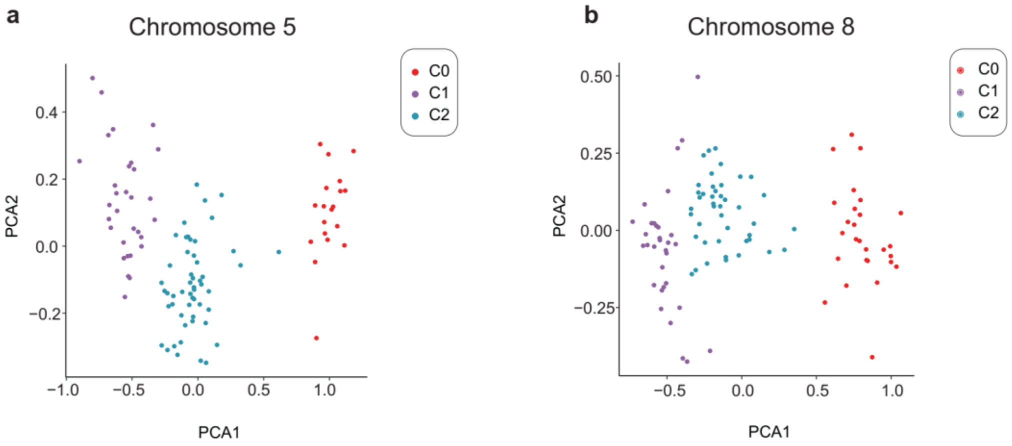


**Supplementary figure S9. The PCA results of sample clustering using n-numbers of HORs in chr5 (a) and chr8 (b).**

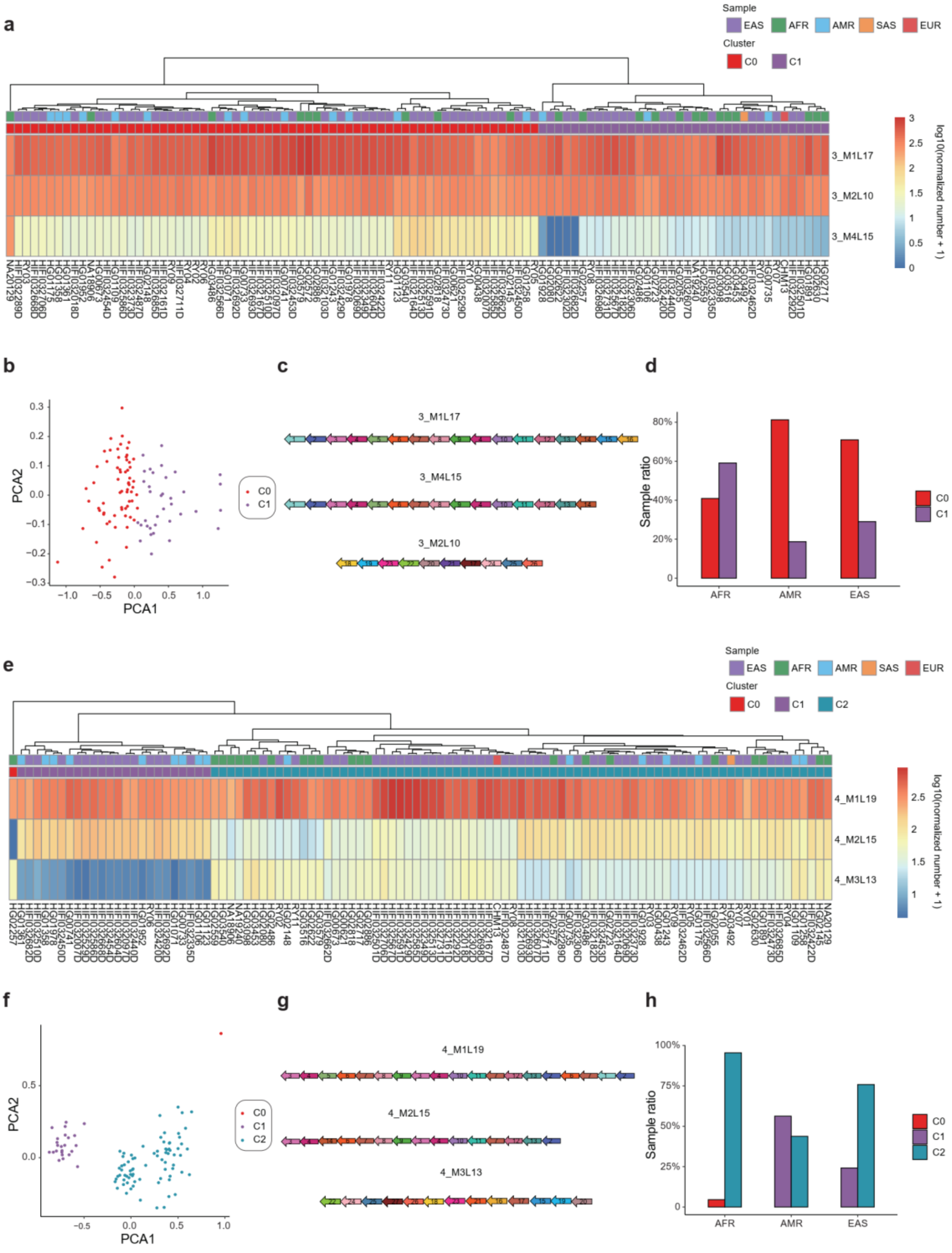


**Supplementary figure S10. Sample clustering based on HORs in chr3 and chr4. a.** The heatmap and sample hierarchical clustering of HOR n-numbers in chromosome 3. **b.** The PCA result of sample clustering using HOR n-numbers in chr3. **c.** Monomer patterns of 3_M1L17, 3_M4L15 and 3_M2L10. **d.** The proportion of samples in each of the AFR, AMR and EAS populations containing 3_C0, 3_C1 and 3_C2. **e.** The heatmap and sample hierarchical clustering of HOR n-numbers in chromosome 4. **f.** The PCA result of sample clustering using HOR n-numbers in chr4. **g.** Monomer patterns of 4_M1L19, 4_M2L15 and 4_M3L13. **h.** The proportion of samples in each of the AFR, AMR and EAS populations containing 4_C0, 4_C1 and 4_C2.

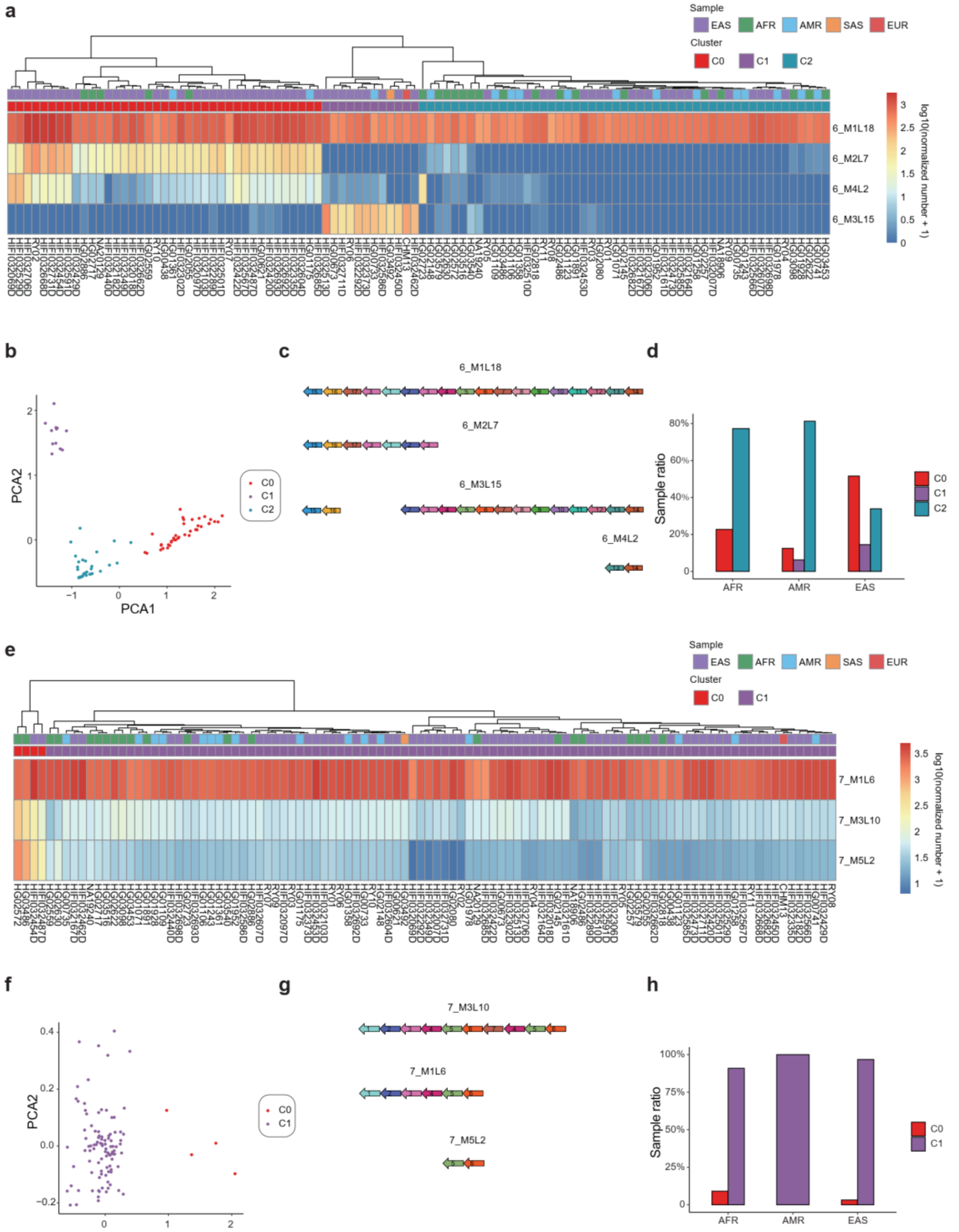


**Supplementary figure S11. Sample clustering based on HORs in chr6 and chr7. a.** The heatmap and sample hierarchical clustering of HOR n-numbers in chromosome 6. **b.** The PCA result of sample clustering using HOR n-numbers in chr6. **c.** Monomer patterns of 6_M1L18, 6_M2L7, 6_M3L15 and 6_M4L2. **d.** The proportion of samples in each of the AFR, AMR and EAS populations containing 6_C0, 6_C1 and 6_C2. **e.**

The heatmap and sample hierarchical clustering of HOR n-numbers in chromosome 7. **f.** The PCA result of sample clustering using HOR n-numbers in chr7. **g.** Monomer patterns of 7_M1L6, 7_M3L10 and 7_M5L2. **h.** The proportion of samples in each of the AFR, AMR and EAS populations containing 7_C0 and 7_C1.

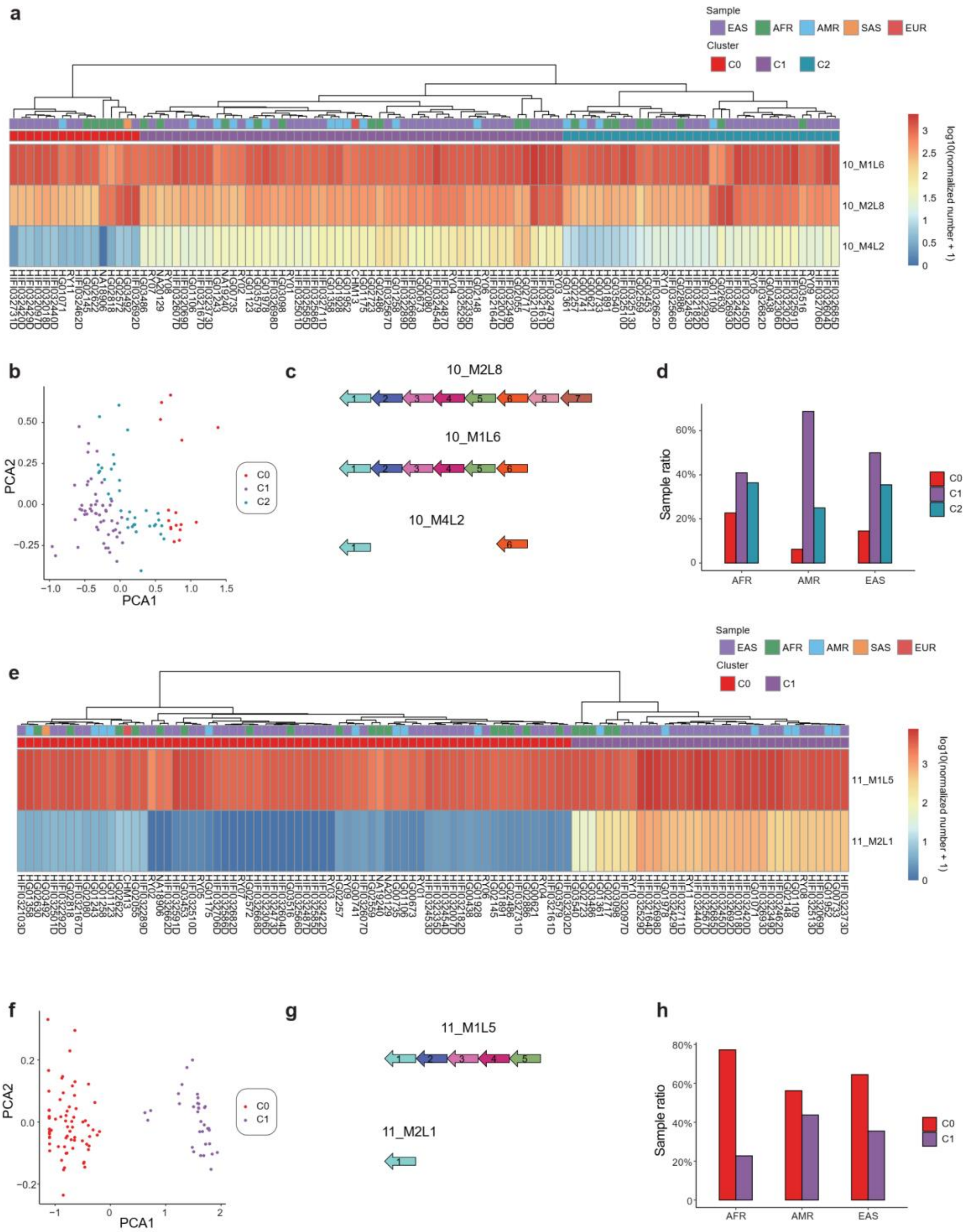


**Supplementary figure S12. Sample clustering based on HORs in chr10 and chr11.** **a.** The heatmap and sample hierarchical clustering of HOR n-numbers in chromosome 10. **b.** The PCA result of sample clustering using n-numbers of HORs in chr10. **c.** Monomer patterns of 10_M1L6, 10_M2L8 and 10_M4L2. **d.** The proportion of samples in each of the AFR, AMR and EAS populations containing 10_C0, 10_C1 and 10_C2.

**e.** The heatmap and sample hierarchical clustering of HOR n-numbers in chromosome 11. **f.** The PCA result of sample clustering using HOR n-numbers in chr11. **g.** Monomer patterns of 11_M1L5 and 11_M2L1. **h.** The proportion of samples in each of the AFR, AMR and EAS populations containing 11_C0 and 11_C1.

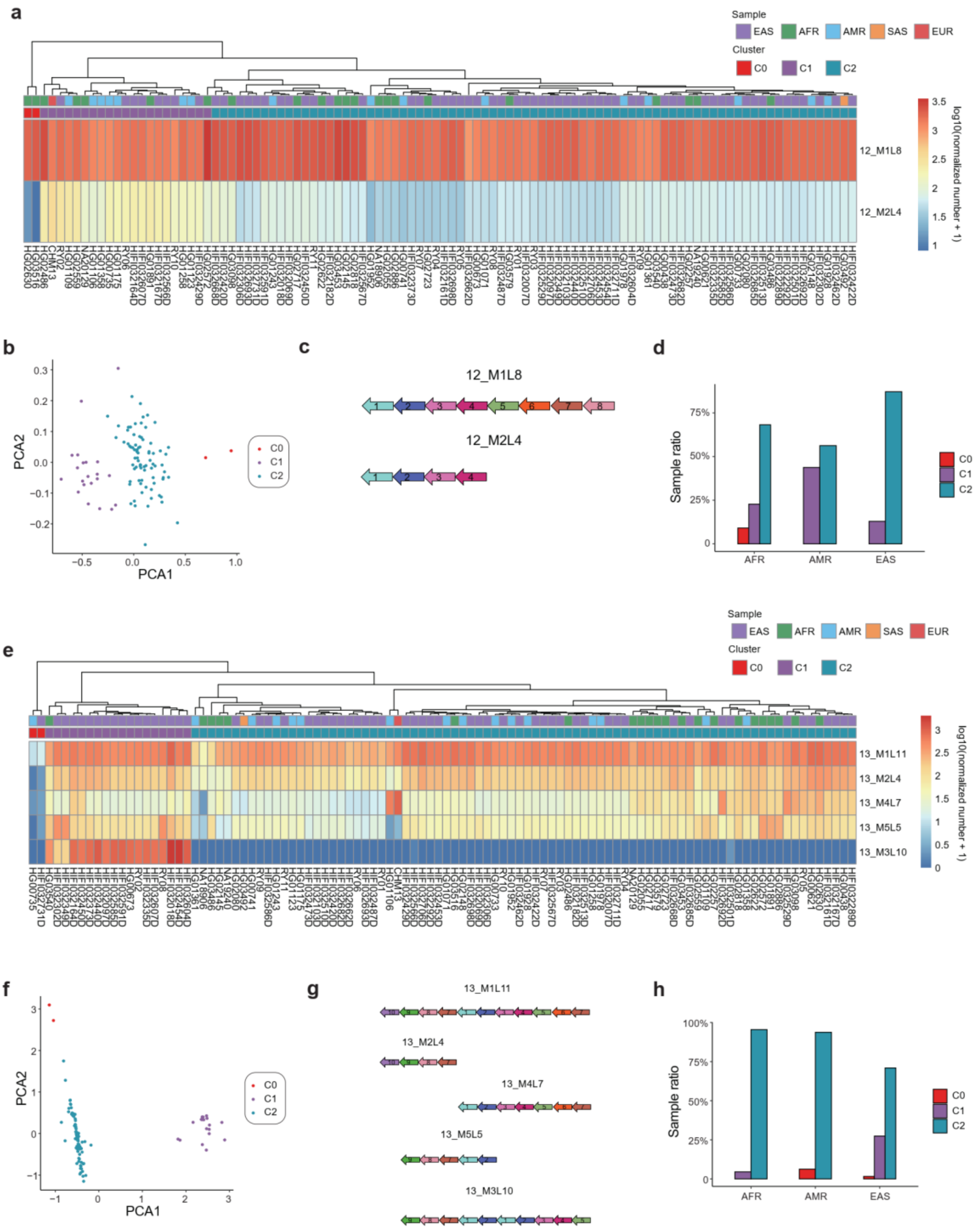


**Supplementary figure S13. Sample clustering based on HORs in chr12 and chr13.** **a.** The heatmap and sample hierarchical clustering of HOR n-numbers in chromosome 12. **b.** The PCA result of sample clustering using HOR n-numbers in chr12. **c.** Monomer patterns of 12_M1L8 and 12_M2L4. **d.** The proportion of samples in each of the AFR, AMR and EAS populations containing 12_C0, 12_C1 and 12_C2. **e.** The heatmap and sample hierarchical clustering of HOR n-numbers in chromosome 13. **f.** The PCA result of sample clustering using HOR n-numbers in chr13. **g.** Monomer patterns of 13_M1L11, 13_M2L4, 13_M4L7, 13_M5L5 and 13_M3L10. **h.** The proportion of samples in each of the AFR, AMR and EAS populations containing 13_C0, 13_C1 and 13_C2.

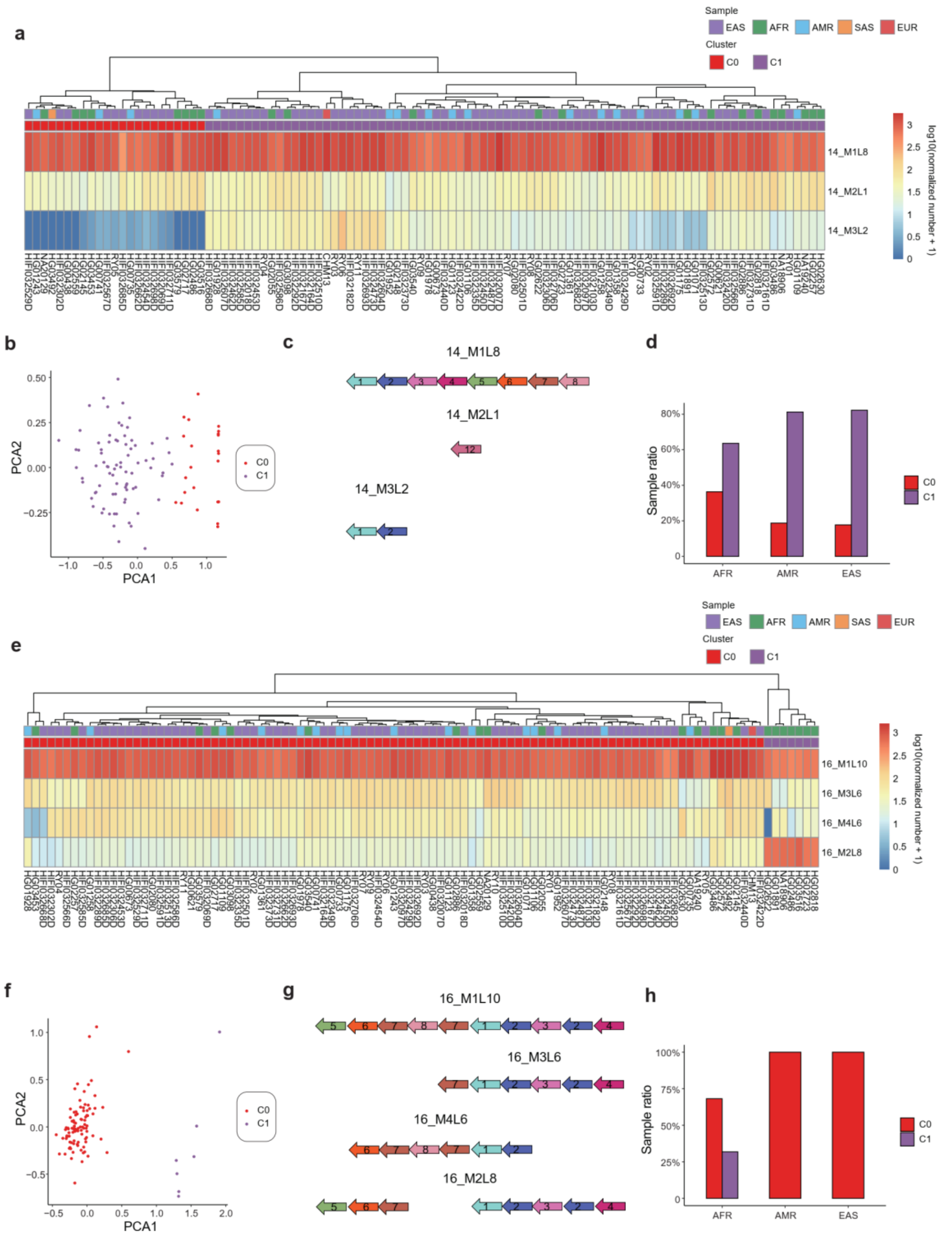


**Supplementary figure S14. Sample clustering based on HORs in chr14 and chr16. a.** The heatmap and sample hierarchical clustering of HOR n-numbers in chromosome 14. **b.** The PCA result of sample clustering using HOR n-numbers in chr14. **c.** Monomer patterns of 14_M1L8, 14_M2L1 and 14_M3L2. **d.** The proportion of samples in each of the AFR, AMR and EAS populations containing 14_C0 and 14_C1. **e.** The heatmap and sample hierarchical clustering of HOR n-numbers in chromosome 16. **f.** The PCA result of sample clustering using HOR n-numbers in chr16. **g.** Monomer patterns of 16_M1L10, 16_M3L6 and 16_M4L6, 16_M2L8. **h.** The proportion of samples in each of the AFR, AMR and EAS populations containing 16_C0 and 16_C1.

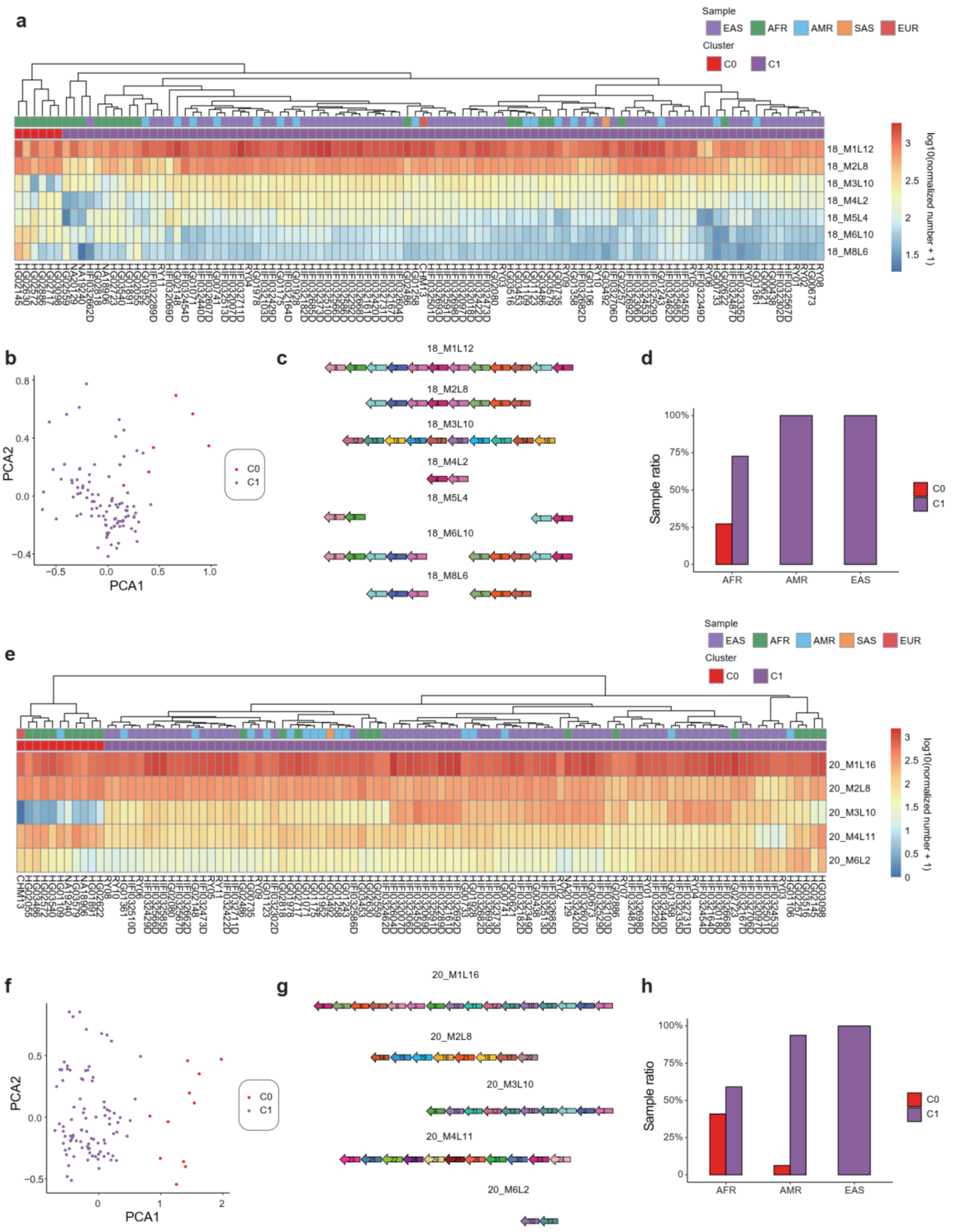


**Supplementary figure S15. Sample clustering based on HORs in chr18 and chr20.** **a.** The heatmap and sample hierarchical clustering of HOR n-numbers in chromosome 18. **b.** The PCA result of sample clustering using HOR n-numbers in chr18. **c.** Monomer patterns of 18_M1L12, 18_M2L6, 18_M3L10, 18_M4L2, 18_M5L4, 18_M6L10 and 18_M8L6. **d.** The proportion of samples in each of the AFR, AMR and EAS populations containing 18_C0 and 18_C1. **e.** The heatmap and sample hierarchical clustering of HOR n-numbers in chromosome 20. **f.** The PCA result of sample clustering using HOR n-numbers in chr20. **g.** Monomer patterns of 20_M1L16, 20_M2L8, 20_M3L10, 20_M4L11 and 20_M6L2. **h.** The proportion of samples in each of the AFR, AMR and EAS populations containing 20_C0 and 20_C1.

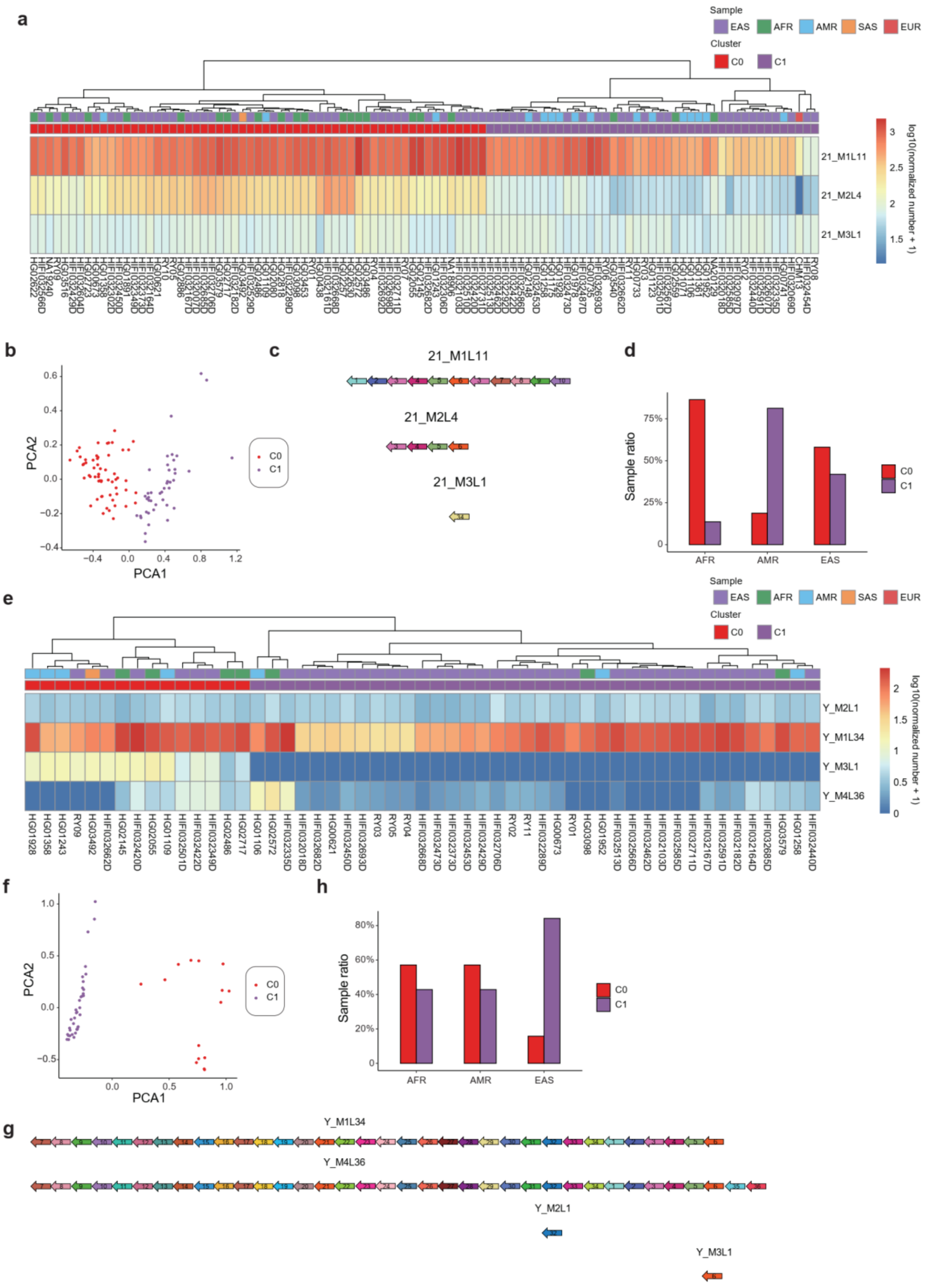


**Supplementary figure S16. Sample clustering based on HORs in chr21 and chrY.**

**a.** The heatmap and sample hierarchical clustering of HOR n-numbers in chromosome 21. **b.** The PCA result of sample clustering using HOR n-numbers in chr21. **c.** Monomer patterns of 21_M1L11, 21_M2L4 and 21_M3L1. **d.** The proportion of samples in each of the AFR, AMR and EAS populations containing 21_C0 and 21_C1. **e.** The heatmap and sample hierarchical clustering of HOR n-numbers in chromosome Y. **f.** The PCA result of sample clustering using HOR n-numbers in chrY. **g.** Monomer patterns of Y_M2L1, Y_M1L34, Y_M3L1 and Y_M4L36. **h.** The proportion of samples in each of the AFR, AMR and EAS populations containing Y_C0 and Y_C1.

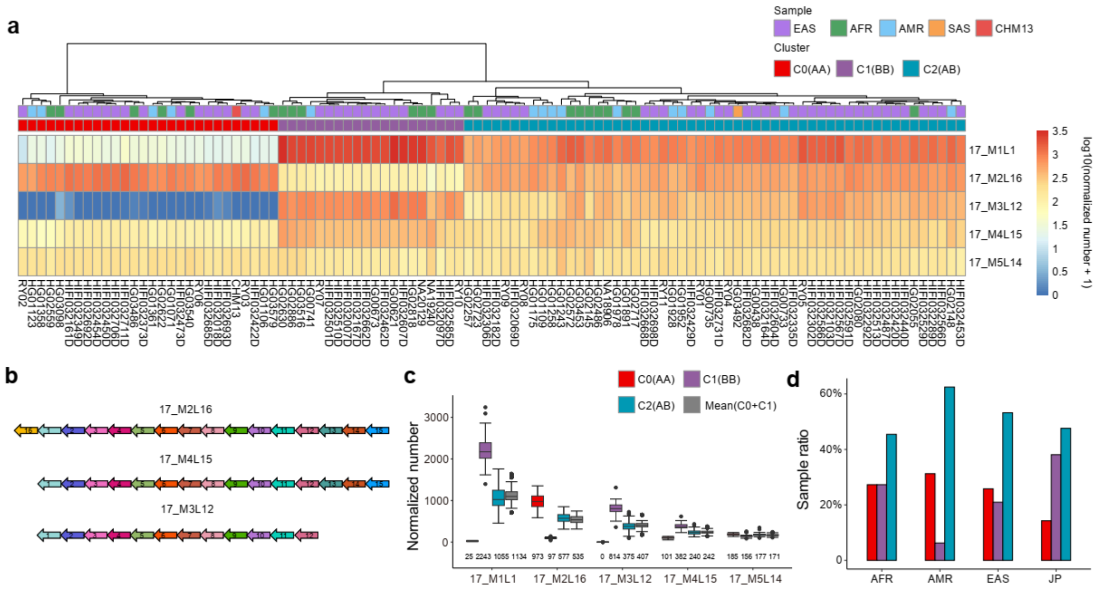


**Supplementary figure S17. Centromere genotype from sample clustering based on HOR n-numbers in chromosome 17. a.** The heatmap and sample hierarchical clustering of HOR n-numbers in chromosome 17. **b.** Monomer patterns of 17_M2L16, 17_M4L15 and 17_M3L12. **c.** The box plot of HOR n-numbers in 17_C0 (AA), 17_C1 (BB) and 17_C2 (AB). For each HOR, the Mean(C0+C1) represents the pairwise mean n-numbers in 17_C0 and 17_C1. **d.** The proportion of samples in each of the AFR, AMR and EAS populations containing 17_C0, 17_C1 and 17_C2. JP is Japanese population.

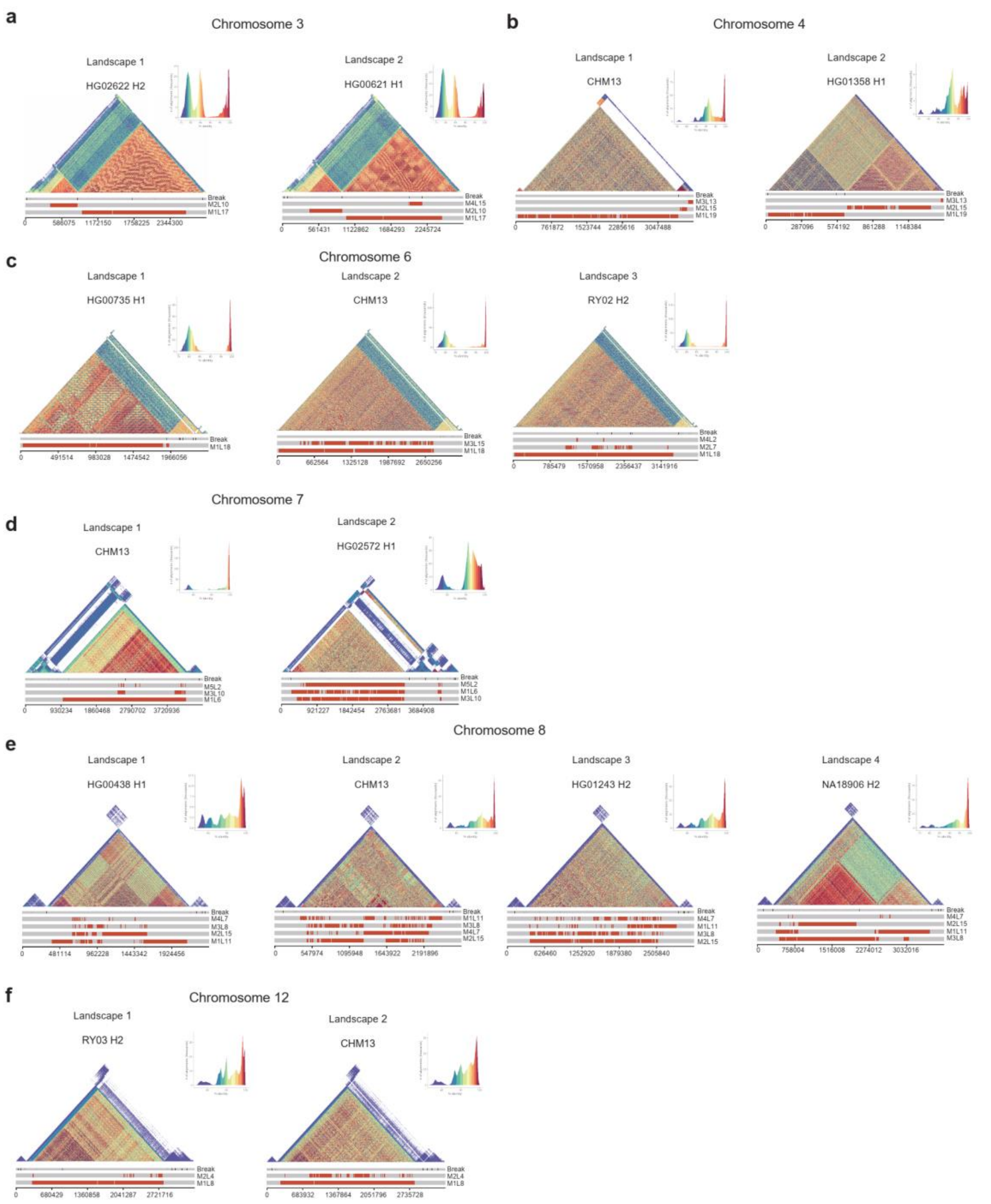


**Supplementary figure S18. HOR landscapes on chromosome 3, 4, 6, 7, 8 and 12. a-f.** The HOR landscapes on chr3 represented by HG02622 H2 and HG00621 H1 (**a**), chr4 represented by CHM13 and HG01358 H1 (**b**), chr6 represented by HG00735 H1, CHM13 and RY02 H2 (**c**), chr7 represented by CHM13 and HG02572 H1 (**d**), chr8 represented by HG00438 H1, CHM13, HG01243 H2 and NA18906 H2 (**e**), chr12 represented by RY03 H2 and CHM13 (**f**). The triangle similarity heatmaps are generated by StainedGlass [3].

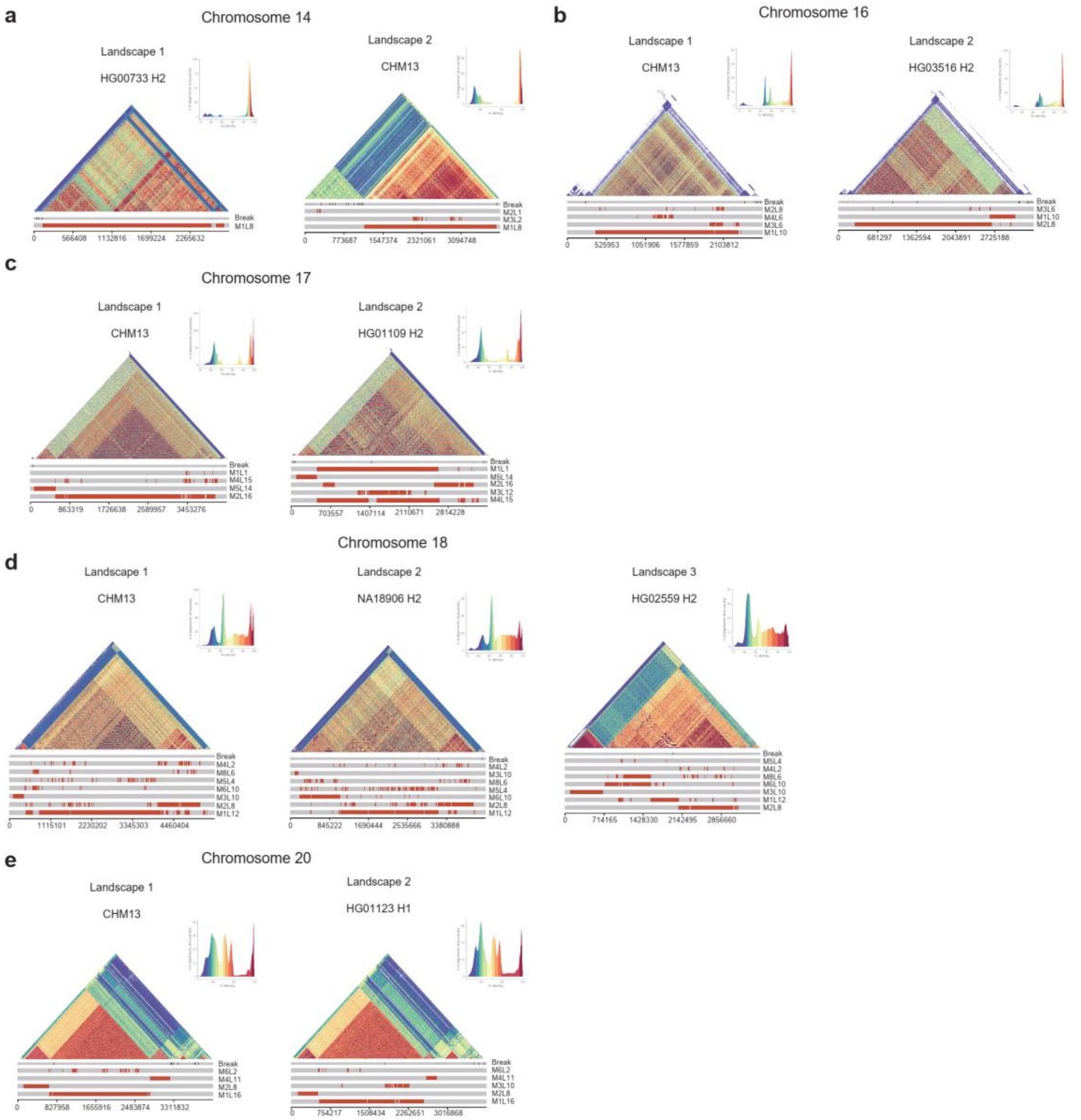


**Supplementary figure S19. HOR landscapes on chromosome 14, 16, 17, 18 and 20.**
**a-e.** The HOR landscapes on chr14 represented by HG00733 H2 and CHM13 (**a**), chr16 represented by CHM13 and HG03516 H2 (**b**), chr17 represented by CHM13 anHG01109 H2 (**c**), chr18 represented by CHM13, NA18906 H2 and HG02559 H2 (**d**), chr20 represented by CHM13 and HG01123 H2 (**e**). The triangle similarity heatmaps are generated by StainedGlass [3].

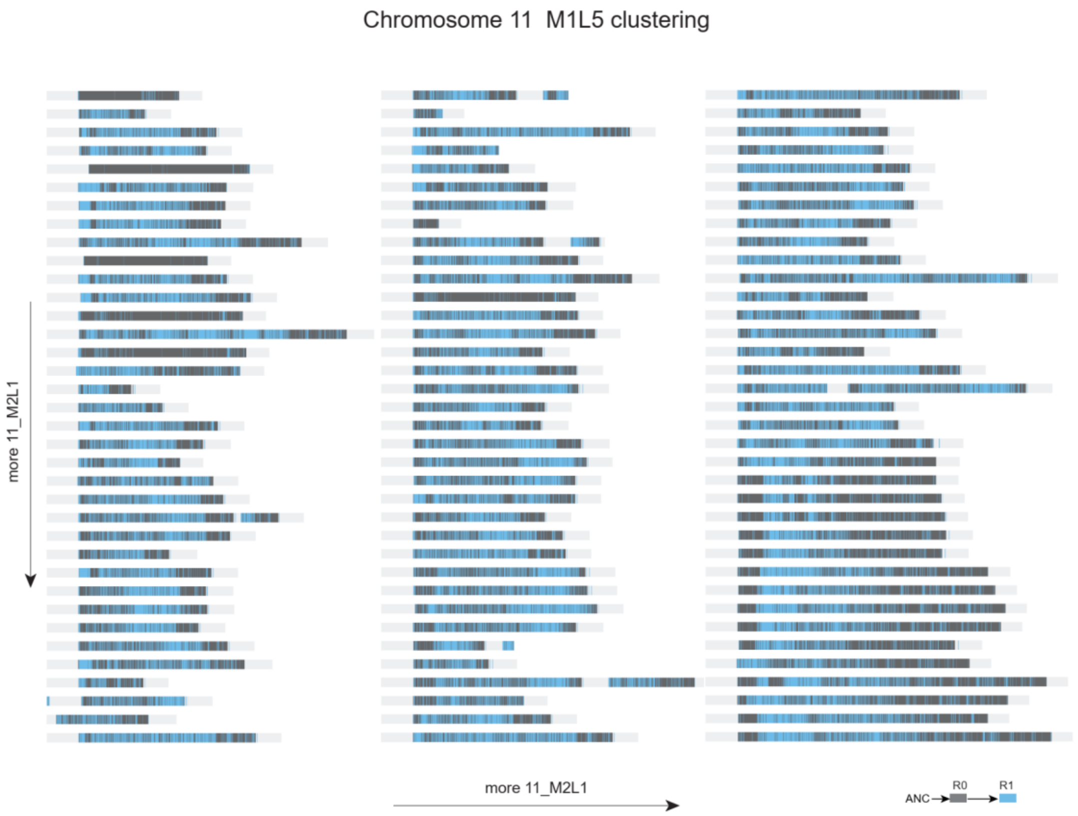


**Supplementary figure S20. Cross-landscape 11_M1L5 clustering of chromosome 11 CASAs for all samples.** Each track is the clustering result of a sample. The samples in each column have increasing number of 11_M2L1 from top to bottom and the content of 11_M2L1 increases between columns from left to right.

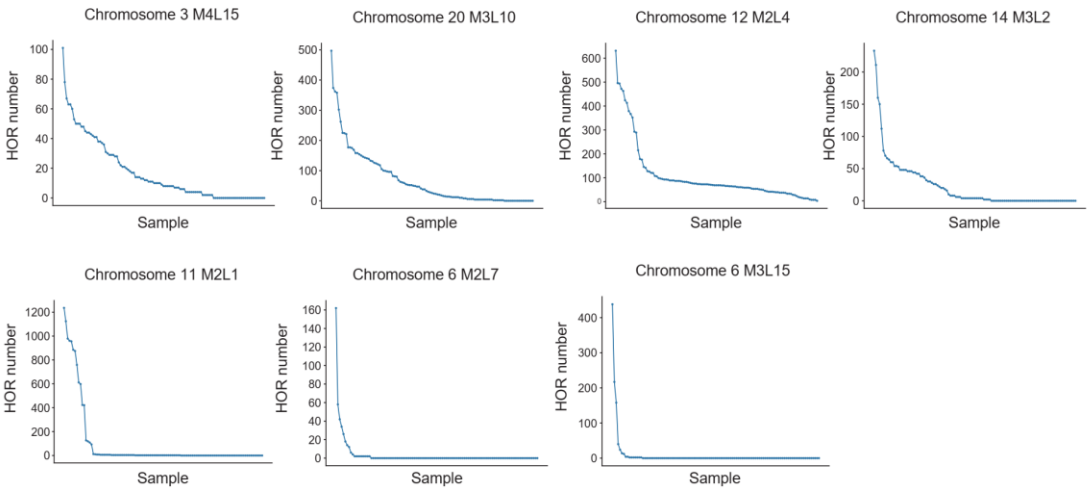


**Supplementary figure S21. LN-HOR number among samples.** The polylines show LN-HOR number in different samples sorted from high to low in chr3, 20, 12, 14, 11 and 6. The local expansion rates differ greatly between chromosomes, appearing relatively high for chr11 but lower for chr3 and 20.

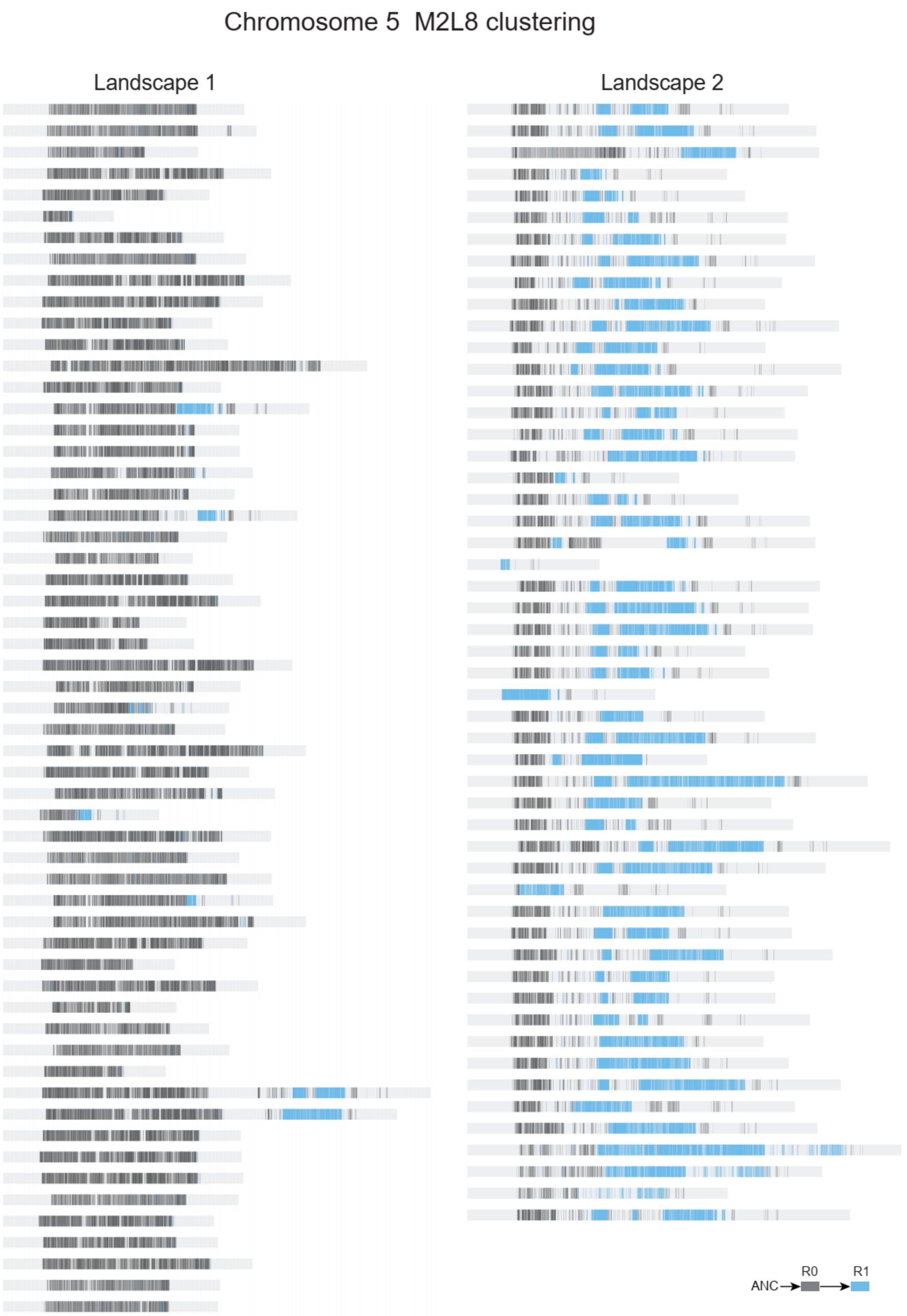


**Supplementary figure S22. Cross-landscape 5_M2L8 clustering result of chr5 CASAs for all samples.** Each track is the clustering result of a sample. Most samples in landscape 1 is mainly composed by R0, and most samples in landscape 2 contains R1 in the middle and right of the CASA.

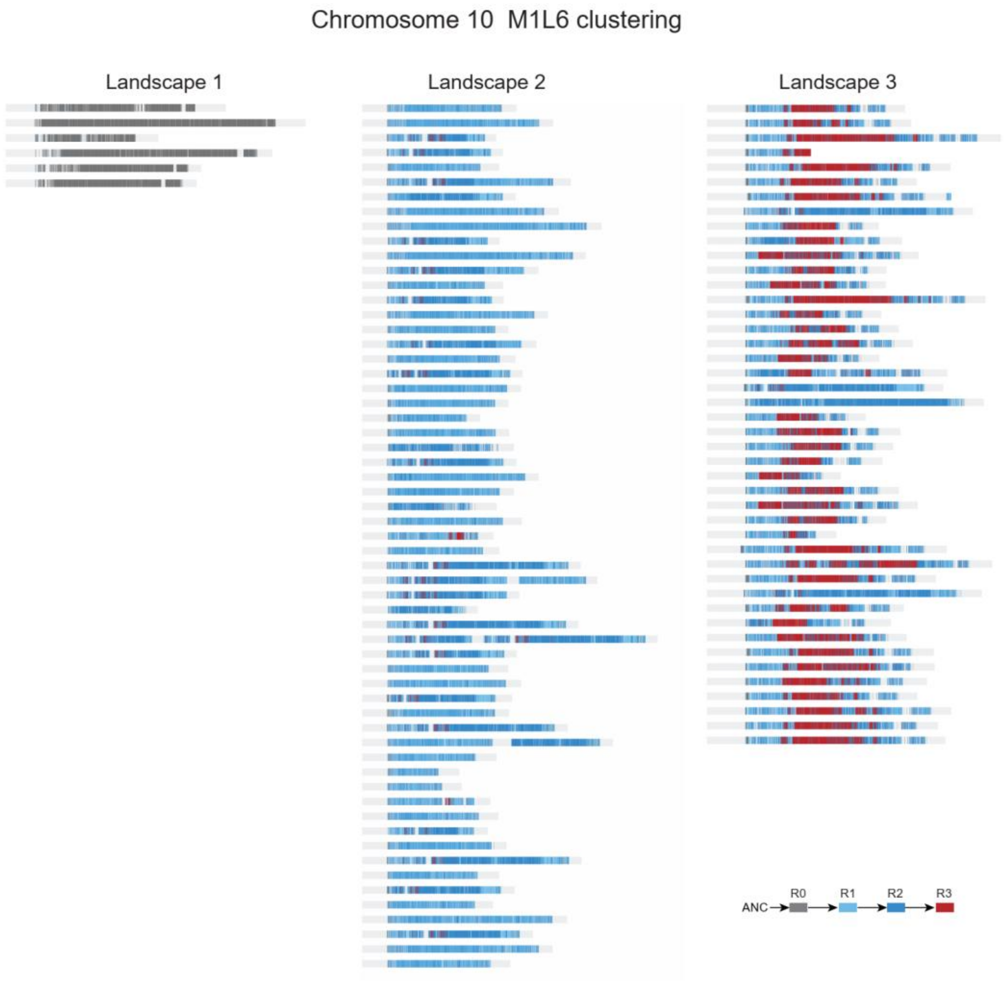


**Supplementary figure S23. Cross-landscape 10_M1L6 cluster result of chr10 CASAs for all samples.** Each track is the clustering result of a sample. Most samples in landscape 1 is mainly composed by R0. In most landscape 2 CASAs, there is a little R0 region on the edge of the HOR region and most of the array is composed by mixed R1 and R2. And in most landscape 3 CASAs, there is R3 in the middle of the array.

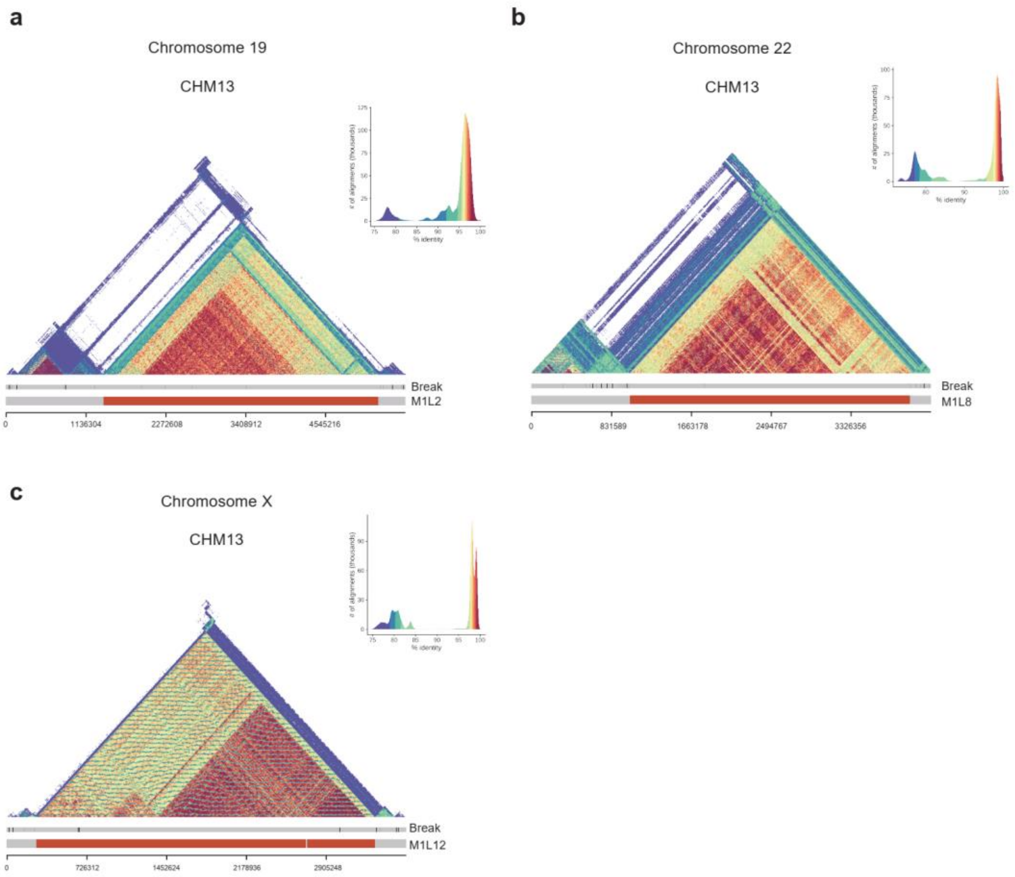


**Supplementary figure S24. HOR landscapes on chromosome 19, 22 and X. a-c.** The HOR landscapes on chr19 (**a**), chr22 (**b**) and chrX (**c**) all represented by CHM13. All the three chromosomes have one homogenous landscape which contains only one primary HOR. The triangle similarity heatmaps are generated by StainedGlass [3].

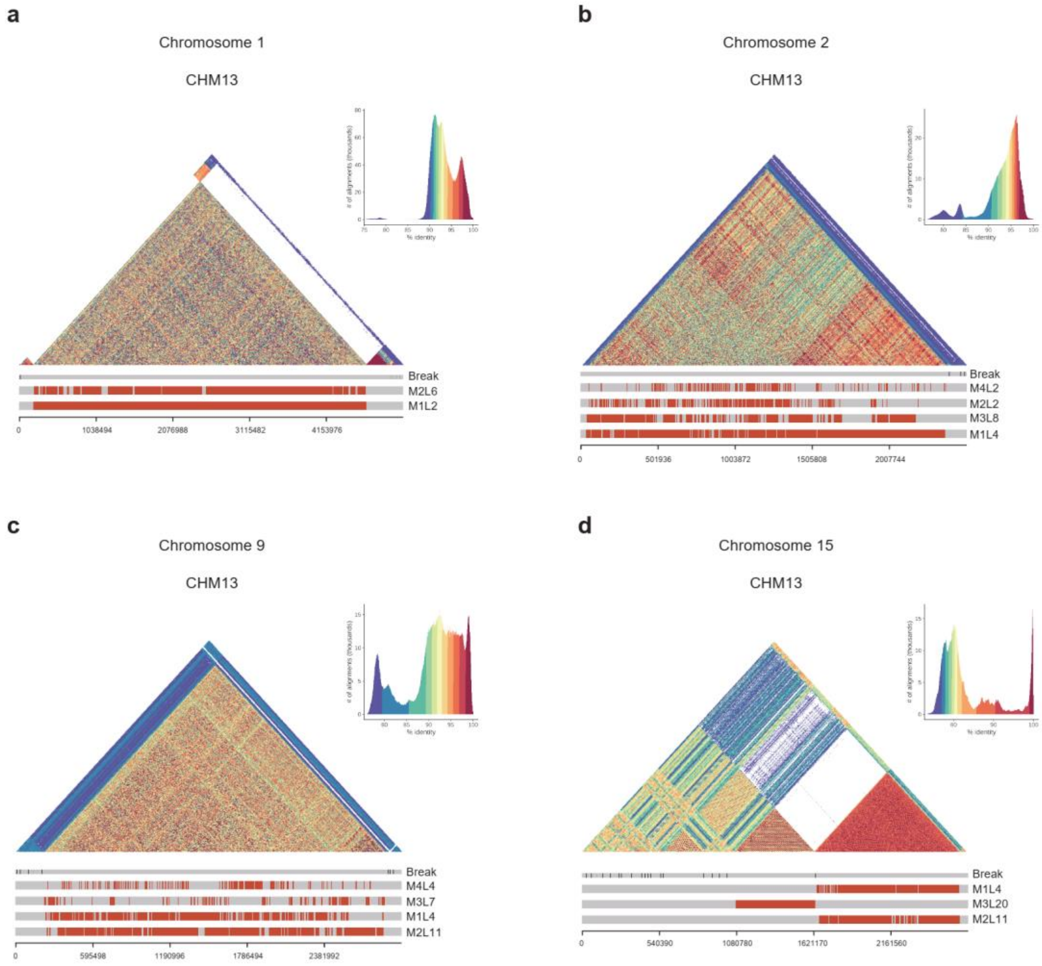


**Supplementary figure S25. HOR landscapes on chromosome 1, 2, 9 and 15. a-d.** The HOR landscapes on chr1 (**a**), chr2 (**b**), chr9 (**c**) and chr15 (**d**) all represented by CHM13. All the four chromosomes have one landscape which contains a large number of LN-HORs. The triangle similarity heatmaps are generated by StainedGlass [3].

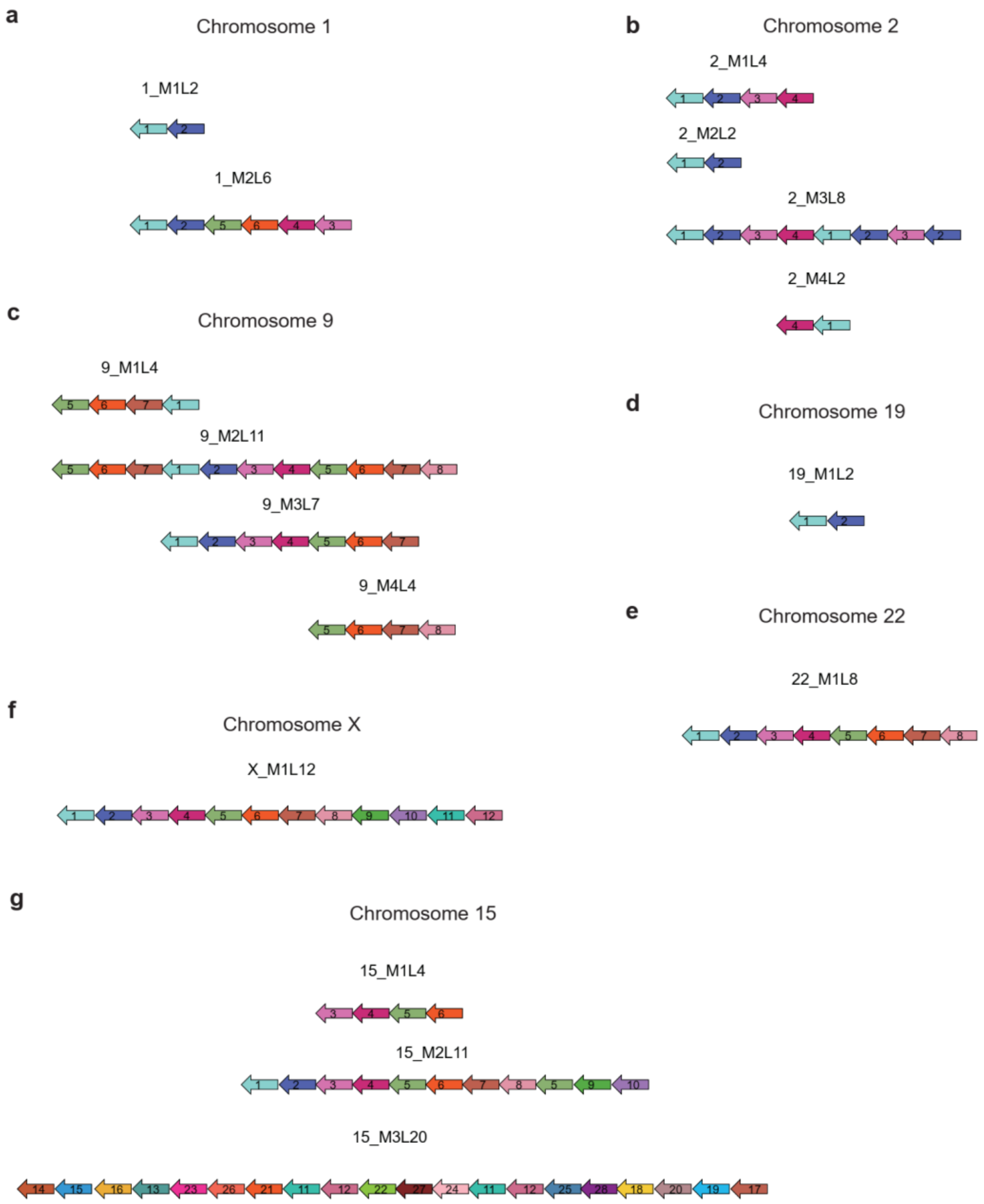


**Supplementary figure S26. Monomer sequences of HORs on chromosome 1, 2, 9, 19, 22, X and 15.**

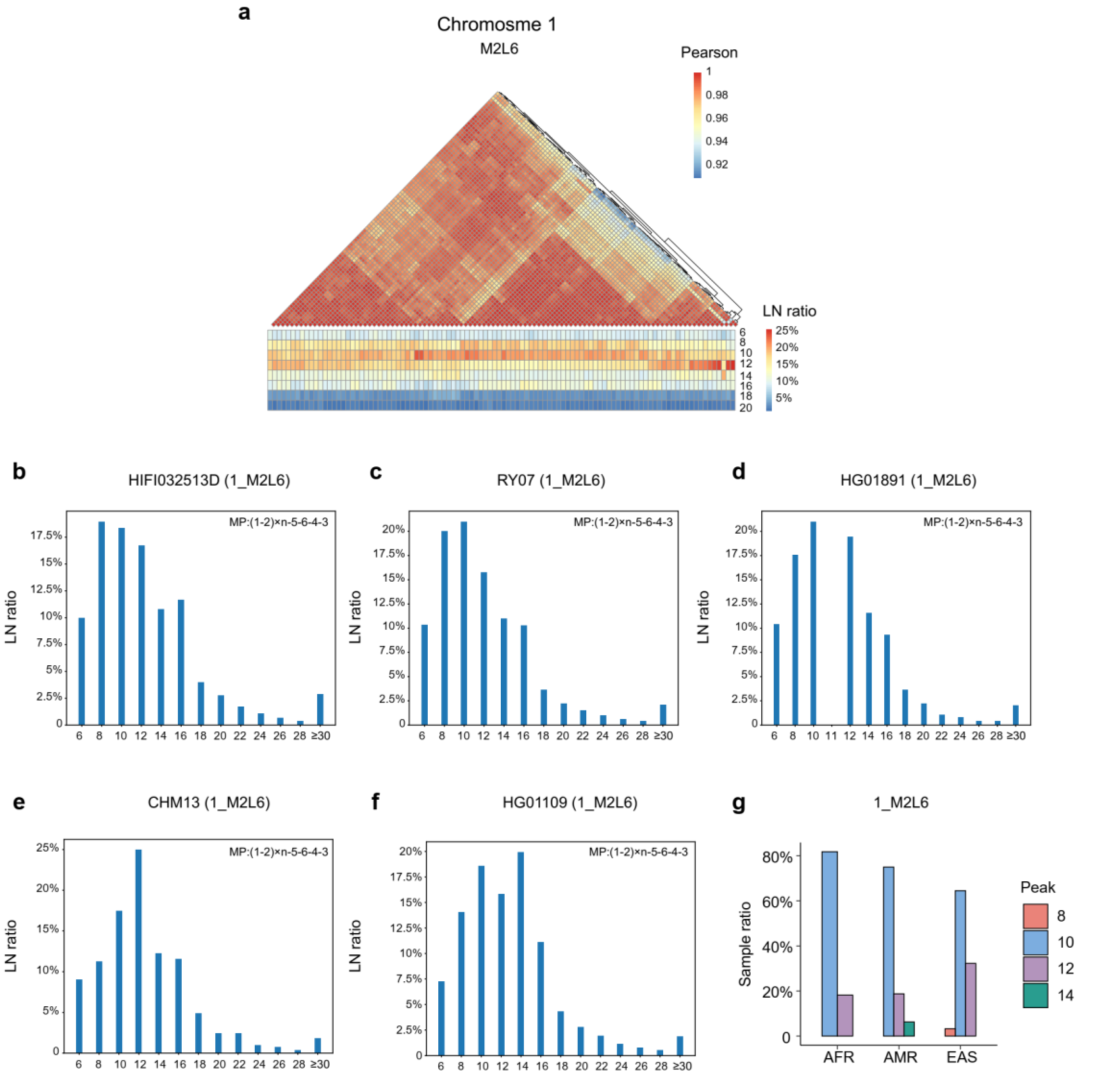


**Supplementary figure S27. 1_M2L6 monomer pattern distribution in different samples. a.** Sample clustering based on Pearson correlation coefficient of 1_M2L6 monomer pattern length. LN ratio is the number of local nested unit compared with the total number of 1_M2L6. **b-f.** The 1_M2L6 unit monomer length distribution of samples with 8-mer peak represented by HIFI032513D (**a**), 10-mer peak represented by RY07 (**b**) and HG01891 (**c**), 12-mer peak represented by CHM13 (**d**) and 14-mer peak represented by HG01109 (**e**). **g.** The sample ratio of 8-mer peak, 10-mer peak, 12-mer peak and 14-mer peak samples among AFR, AMR and EAS populations.

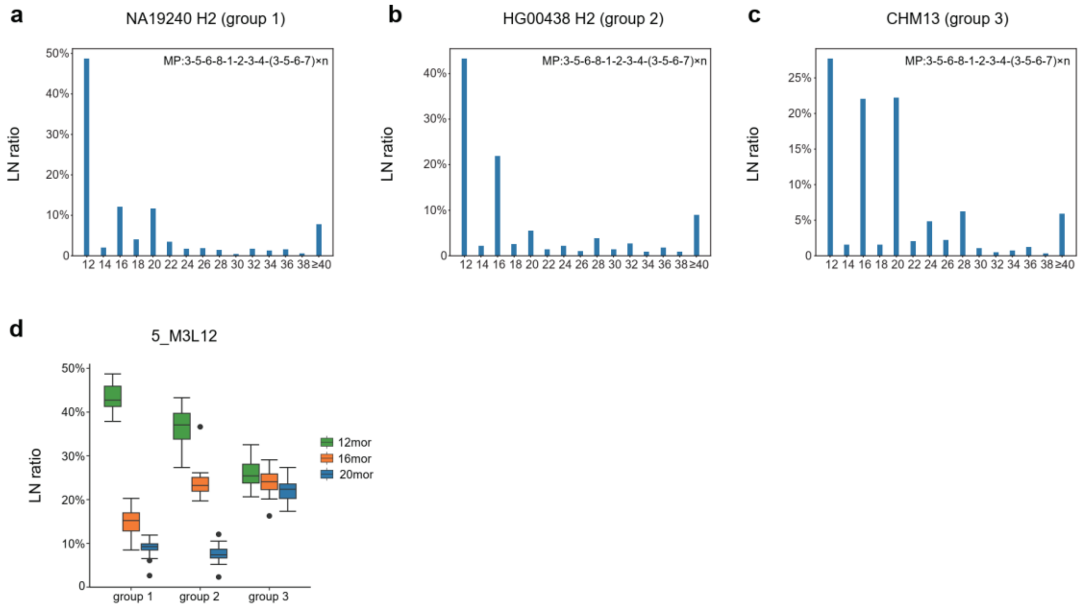


**Supplementary figure S28. 5_M3L12 monomer pattern distribution in different groups. a-c.** The 5_M3L12 unit monomer length distribution of samples in different groups. Samples in group 1 are represented by NA19240 H2 (**a**). Samples in group 2 are represented by HG00438 H2 (**b**). Samples in group 3 are represented by CHM13 (**c**). **d.** The 12mor, 16mor and 20mor M3L12 distribution of samples in group 1, 2 and 3.

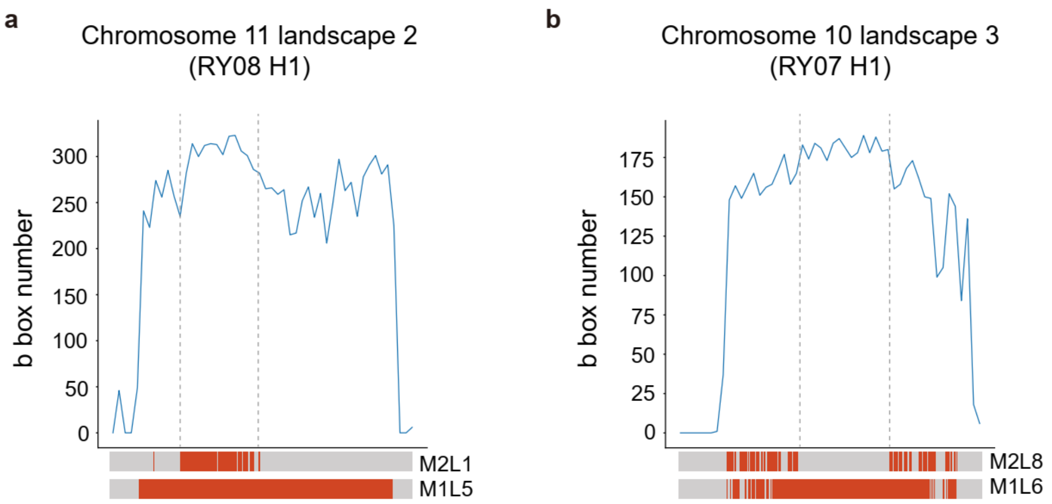


**Supplementary figure S29. b box density in different regions of the chromosome.** **a.** The b box density of RY08 H1 chr11 centromere array. The M2L1 local nested region shows higher b box density than flanking regions. **b.** The b box density of RY07 H1 chr10 centromere array. The medium region with homogeneous M1L6 HORs shows higher b box density than flanking regions. The density window number is 50.

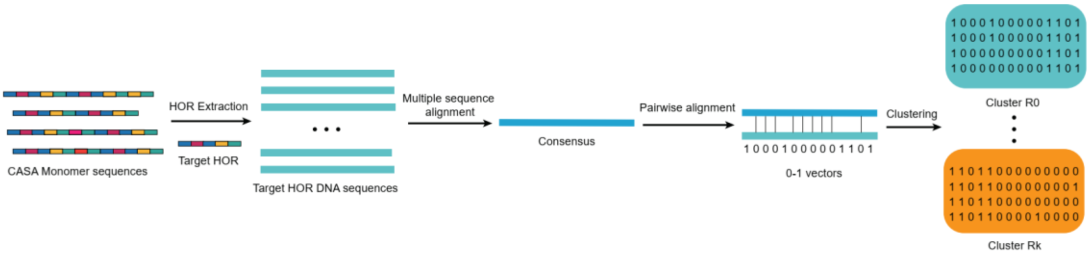


**Supplementary figure S30. Cross-landscape HOR clustering workflow.** Target HOR DNA sequences for clustering are first extracted from CASA monomer sequences based on HiCAT-human-assembly annotation. Then, multiple sequence alignment is performed based on HOR DNA sequences to generate HOR consensus sequence. Each HOR DNA sequence is compared with consensus to obtain 0-1 vector, where 0 indicates this HOR shares the same base with consensus sequence at that position and 1 indicates there is a difference. Finally, the 0-1 vectors of all HOR unit are clustered based on k-means. For each target HOR, we chose the smallest k that can show the difference among landscapes.

### Supplementary tables

**Supplementary table S1.** Accession numbers and sample metadata for HiFi reads and assemblies

**Supplementary table S2.** Reads classification recall in 30X HiFi simulation test

**Supplementary table S3.** HOR cover ratio in error classified reads for all alpha satellite regions

**Supplementary table S4.** The estimate HOR array size

**Supplementary table S5.** The normalized number of HORs among samples in autosomes.

**Supplementary table S6.** The normalized number of HORs among samples in chromosome X.

**Supplementary table S7.** The normalized number of HORs among samples in chromosome Y.

**Supplementary table S8.** The mean fold-change of HORs among samples in autosomes.

**Supplementary table S9.** The mean fold-change of HORs among samples in chromosome X.

**Supplementary table S10.** The mean fold-change of HORs among samples in chromosome Y.

**Supplementary table S11.** The standard deviation of the mean fold-change of HORs among samples

**Supplementary table S12.** The normalized numbers of 33 v-HORs among different populations

**Supplementary table S13.** Spearman correlation between all HORs.

**Supplementary table S14.** Sample clustering based on HORs on chromosome with v-HORs

**Supplementary table S15.** HOR normalized number comparison of different clusters in HiFi reads data on chromosome 5, 8 and 17

**Supplementary table S16.** Identity between reconstructed ancestral HOR sequence and consensus HOR sequence in each cluster

**Supplementary table S17.** Monomer pattern sample distribution in chromosome 1 M2L6

**Supplementary table S18.** The 12mor, 16mor and 20mor M3L12 distribution of samples in group 1, group 2 and group 3

**Reference**


1. Wenger, A.M. *et al.* Accurate circular consensus long-read sequencing improves variant detection and assembly of a human genome. *Nat Biotechnol* **37**, 1155-1162 (2019).
2. Altemose, N. *et al.* Complete genomic and epigenetic maps of human centromeres. *Science* **376**, eabl4178 (2022).
3. Vollger, M.R., Kerpedjiev, P., Phillippy, A.M. & Eichler, E.E. StainedGlass: interactive visualization of massive tandem repeat structures with identity heatmaps. *Bioinformatics* **38**, 2049-2051 (2022).
